# Mitochondria-Targeted Peptides and Bilayer Composition Modulate Membrane Electroporation Under Elevated Electrochemical Stress

**DOI:** 10.1101/2025.08.05.668743

**Authors:** Jeffrey D. Tamucci, Adam Zweifach, Nathan N. Alder, Eric R. May

**Affiliations:** Department of Molecular and Cell Biology, University of Connecticut, Storrs, CT, 06269, U.S.A

## Abstract

Several studies have used molecular dynamics (MD) simulations to examine the relationship between transmembrane potentials *(ΔΨ_m_*) and electroporation; however, research on how this relationship presents in complex membranes with heterogenous lipid compositions is limited. Here we use all-atom, double-bilayer MD simulations with explicitly-modeled *ΔΨ_m_* voltages to compare how membranes with homogenous vs. heterogenous lipid compositions respond to varying degrees of electrochemical stress. These two bilayer systems were also exposed to a mitochondria-targeted peptide named Elamipretide which has been shown experimentally to modulate membrane electrostatics. Additionally, we used Computational Electrophysiology (CompEL) MD simulations to analyze how lipid composition and peptide exposure influence rates of passive transmembrane ion flux during electroporation events. The addition of cardiolipin (CL) and Elamipretide increased membrane capacitance, decreasing the *ΔΨ_m_* for a given transmembrane charge imbalance (*δ_TM_*) and protecting bilayers against electroporation. These results expand our understanding of how more complex, biologically-relevant bilayers respond to electrochemical stress.

## Introduction

Biomembranes generate complex electrostatic environments in their vicinity, which are critical to mediating their interactions with proteins, ions and other molecular entities. These interactions are essential for cellular processes, ranging from signal transduction to maintaining homeostasis. In certain biological environments, membranes are exposed to transmembrane potentials which cells utilize in multiple ways including transmembrane transport and energy storage. At the inner mitochondrial membrane (IMM), the electrochemical proton potential 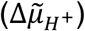 is the primary energy source that drives ATP synthesis and other energy-related processes.^1^ Understanding the factors that influence the generation and response to different magnitudes of transmembrane potentials, particularly the influence of membrane composition and membrane-protein/peptide interactions, is of significant biological and therapeutic interest.

The 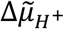 is composed of the proton concentration difference (Δ*pH*) and the electrostatic potential difference (*ΔΨ_m_*), which are coupled and typically equate to ∼0.5 pH units and ∼150-180 mV across the IMM in healthy mitochondria.^1^ Mitochondrial dysfunction is often associated with decreased Δψ_m_ (depolarized IMM), as occurs with impaired electron transport complex activity or extensive proton leak across the IMM.^2,3^ On the other hand, some pathological conditions can cause excessively high *ΔΨ_m_* (hyperpolarized IMM), including cancer,^4^ abnormal T cell activation,^5^ defects in ATP synthase activity or ADP availability,^6,7^ and fatty acid oxidation disorders.^8^ In other cases such as ischemia-reperfusion injury, the Δψ_m_ shows unstable oscillations between depolarization and hyperpolarization.^9^ A recent study revealed that experimentally increasing the saturation index of mitochondrial phospholipids led to aberrant IMM morphology and ATP synthase activity, concurrent with extreme hyperpolarization.^10^ A known outcome of increased *ΔΨ_m_* is unregulated reactive oxygen species (ROS) production;^11^ however, the structural and functional consequences of these elevated potentials on mitochondrial membranes remain poorly understood.

Our previous studies^12–14^ have focused on a family of synthetic mitochondria-targeted peptides (MTPs, also known as SS-peptides), which accumulate in mitochondria 1000- to 5000-fold (most likely at the IMM) and mitigate mitochondrial dysfunction^15–17^ in a range of common diseases and conditions.^18–27^ We characterized how the MTPs modulate membrane electrostatic properties, notably finding that they partially restore the IMM’s *ΔΨ_m_*in dysfunctional mitochondria. In our most recent study,^14^ we used all-atom molecular dynamics (MD) simulations to investigate the structural mechanisms through which MTPs influence membrane potential (*Ψ*), finding that they alter bilayer dimensions predictive of an increase in the membrane’s capacitance (*C_m_*), among other effects.

MD simulations are a well-established and widely-used method for studying membrane electrostatics.^28–47^ Of particular relevance is the work by Lin & Alexander-Katz, ^29^ in which they employed double membrane coarse-grained simulations of homogenous lipid bilayers and a range of *ΔΨ_m_* strengths (0 to ∼2.5 V). In that work, they describe a relationship between *ΔΨ_m_* and the transmembrane ionic imbalance per lipid (*e*/Lip), delineating a linear, yielding, and threshold regime. The authors use these three regimes to explain how *ΔΨ_m_* scales linearly with *e*/Lip values to a point at which *ΔΨ_m_* plateaus and bilayers begin occasionally yielding to electroporation (∼1.8 V). Beyond the yielding range, membranes reach a threshold *e*/Lip at which electroporation always occur and pores rapidly emerge within 80 ns. By examining various bilayer properties (melting temperature, order parameters, diffusion), they also concluded that high ionic imbalance “softened bilayers,” making them more “fluid-like” and hypothesized that this would considerably increase their susceptibility to pore formation, even at *ΔΨ_m_* voltages below the yielding range in more complex, biological membranes.

In the current study, we extend the work of Lin & Alexander-Katz in several respects. While they provided foundational insights into how *ΔΨ_m_* impacts the biophysical properties and behaviors of simple bilayers, our understanding of how *ΔΨ_m_* voltages influence the dynamics of more realistic membranes with heterogenous lipid mixtures remains limited. In this study, we use all-atom double-bilayer MD simulations as a reductionist model of the IMM, with each bilayer dividing our “mitochondria” (i.e., simulation box) into two solvent compartments, representing the “intermembrane space” (IMS) and the mitochondrial “matrix.” We examine the effects of *ΔΨ_m_* on membranes composed of pure POPC (palmitoyloleoyl-phosphatidylcholine) and mixtures of POPC and tetraoleoyl-cardiolipin (CL). CL is a unique lipid found almost exclusively in the mitochondria in eukaryotic cells that is critical for optimal assembly and function of IMM protein complexes^10,48,49^ as well as regulation of the mitochondrial *ΔΨ_m_*.^50,51^ Additionally, we investigate the impact of the MTP SS-31 (D-Arg–3,5-dimethyl-Tyr–Lys–Phe-NH_2_) on these bilayers under electrochemical stress. Our simulations reveal that both lipid composition and SS-31 alter the probability of nanopore formation (i.e. electroporation) and spontaneous unassisted ion transport through the lipid bilayer. Additionally, we performed simulations under constant *ΔΨ_m_*, analogous to a voltage-clamp experiments, to investigate the influence of CL and/or SS-31 on the rates of ion leakage. These findings expand upon the work of Lin & Alexander-Katz by incorporating heterogeneous bilayers, MTP interactions, and higher resolution all-atom models, deepening our understanding of how complex lipid bilayers respond to elevated electrochemical stress and the mechanisms by which MTPs and CL can modify the response.

## Methods

### MD Simulations

#### System Preparation

Double-bilayer MD simulations under the influence of a transmembrane potential (*ΔΨ_m_*) were used to characterize the effects of heterogeneous lipid compositions and membrane-interacting peptides on bilayer properties, membrane ion distributions, and transmembrane ion leakage. All-atom single-bilayer systems with explicit membranes and solvent were initially prepared using CHARMM-GUI.^52–55^ Two single-bilayer systems were generated, one with all POPC lipids and the other with POPC and TOCL lipids at a molar ratio of 20:80 TOCL:POPC. A representative double bilayer system is shown in **Fig. 1A** and lipid structures are shown in **Fig. 1B**. Each single bilayer system contained a total of 150 lipids (75 per leaflet). All systems were energy minimized for 5000 steepest descent steps, followed by NVT equilibration for 100 ps with a 1 fs timestep, 200 ps of NPT equilibration with a 1 fs timestep, and ∼100 ns of NPT equilibration with a 2 fs timestep. The semi-isotropic pressure coupling scheme, Berendsen barostat, and Berendsen thermostat were used during NPT equilibration.^56^ All minimization, equilibration, and production simulations were performed using the 2022 version of GROMACS.^57–59^ Position and dihedral restraints were used during equilibration on the lipids to maintain lipid geometry and bilayer morphology.

**Figure 1.**
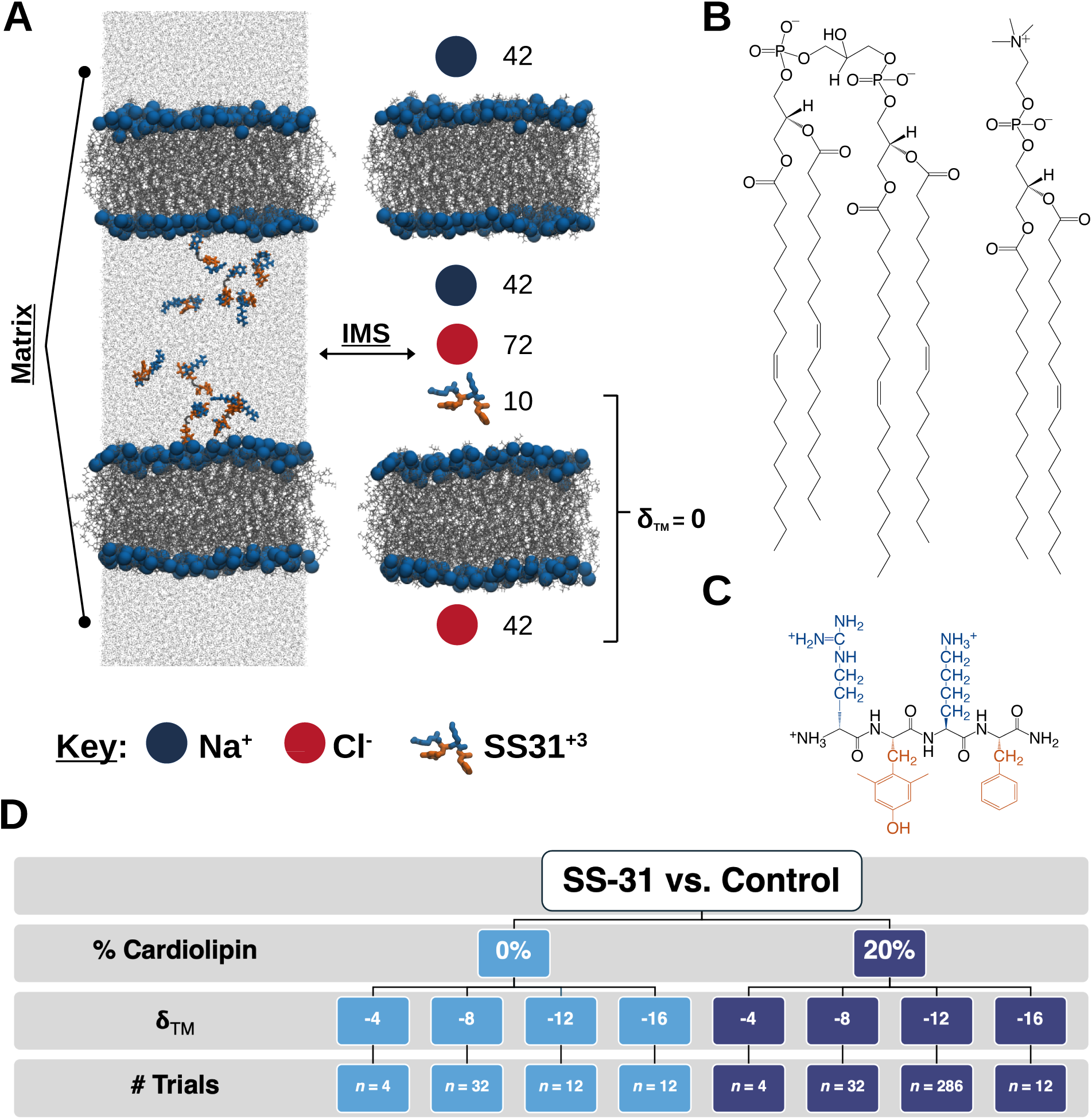
MD system representation, simulation approach, and structures of lipids. (A) System representation and corresponding schematic representation of the initial state of a 20:80 TOCL:POPC peptide-bilayer system without an ion imbalance (𝝳_TM_) between the intermembrane space (IMS) and mitochondrial matrix. Peptides (basic residues = blue; aromatics = orange) suspended in the solvent (gray dots) and the upper and lower membranes (lipid phosphates = blue spheres; lipid acyl chains = gray wireframe) separating the IMS and matrix water compartments. (B) Structures of tetraoleoyl-cardiolipin (TOCL, left) and palmitoyloleoyl-phosphatidylcholine (POPC, right). (C) Structure of SS-31 (also referred to as Elamipretide). (D) Flowchart representation of the unbiased (i.e., non-CompEL, 500 ns/trial) MD simulations run in this study.

The double-bilayer systems were generated by duplicating the single-bilayer systems in the *Z*-direction using the GROMACS *gmx genconf* function.^57,58^ In SS-31-containing systems, 10 peptides were randomly placed in the central solvent compartment at least 1-2 nm from either bilayer’s headgroup region. A description of how the structure and parameters for SS-31 were generated was described in detail previously (see SS-31’s structure in **Fig. 1C**).^12,13^ All systems were re-solvated to ∼70% water by mass and ions were added to a salt concentration to 100 mM NaCl plus neutralizing ions. In membrane-only (“*Apo*”) simulations, an additional 30 Na^+^ ions were added to the central compartment (plus neutralizing ions, if needed) to have a consistent number of positive charges in the *Apo* and SS-31–containing systems. A more in-depth explanation of how one can prepare double-bilayer systems is available here,^60^ and full table schematics of each system in this study are available in the supplement (**Figs. S1-S4**). With peptides position-restrained in the solvent phase, the double-bilayer systems were then minimized and equilibrated following an identical protocol to the one described above except the NPT portion of equilibration was run for 50 ns.

Systems with four different initial *ΔΨ_m_* values were generated in a stepwise fashion by swapping the coordinates of one Cl^-^ and Na^+^ ion in the central (IMS) and outer (Matrix) water compartments respectively to create a transmembrane charge imbalance (*δ_TM_*). After each ion swap, production simulations were performed for multiple trials to equilibrate the membranes and ions. After 75 ns of simulation time, we selected a trial that had not experienced ion leakage and then used that system as a starting point for the next highest *δ_TM_*. Starting with the swap of the second ion pair, the systems were also brought through the CHARMM-GUI default equilibration for bilayers (w/ NPT equilibration for ∼100 ps) before starting their production simulations. Ultimately, we created all PC and 20% CL double bilayer systems in the presence and absence of peptides with one, two, three, and four ions swapped, respectively generating a *δ_TM_* of −4, −8, −12, and −16. Note that we have given *δ_TM_* values a negative sign to align with the convention of describing the IMM’s *ΔΨ_m_* using negative voltage (i.e., −150 mV).

#### Simulation Details

We conducted production simulations probing SS-31’s influence on pore formation and ion leakage in double bilayer systems in two ways: 1) in unbiased ion imbalance simulations where an initial transmembrane potential could be dissipated through pore formation and ion redistribution across the bilayer, and 2) in the presence of a constant *ΔΨ_m_* that persisted even if pores formed. For the unbiased ion imbalance simulations, production runs of 500 ns were initially run for four trials for the systems with an *δ_TM_* of −4 and eight trials for all other *δ_TM_* values.

We aimed to investigate CL’s and SS-31’s influence on lipid bilayers at the lowest magnitude *δ_TM_* in which pores would form (i.e. in the “yielding” regime), which for all PC was *δ_TM_* =-8 and for 20% CL was *δ_TM_* =-12. Based on the proportion of initial trials in which pores formed, we used ClinCalc’s Sample Size Estimator to calculate the number of trials necessary to find a statistically significant effect for a dichotomous endpoint, two independent sample study with an alpha level of 0.05 and statistical power of 0.8.^61^ Accordingly, we ran the estimated number of trials for the yielding range of both lipid compositions (**Fig. 1D**). We also added additional trials to the highest *δ_TM_* systems to increase our sampling before running analyses.

To perform simulations at a constant *ΔΨ_m_*, the GROMACS Computational Electrophysiology (CompEL) module^36,62^ was used to maintain a constant imbalance of Na^+^ and Cl^-^ ions between the IMS and matrix. The CompEL module regularly checks the number of ions in each water compartment. When an ion has crossed from one compartment to the other, it swaps the displaced ion’s coordinates with those of a water molecule in its original compartment, thus maintaining a constant ionic imbalance. A cylinder centered on the bilayer’s center of mass (COM) with a radius of half the diagonal of the bilayer and height of two nm was used to define the channel through which ions could pass. The number of ions in each compartment was checked every 0.2 ps. CompEL simulations were run for both bilayer lipid compositions with and without SS-31 at *δ_TM_* values of −8, - 12 and −16 for the all PC system and −12 and −16 for the 20% CL system. To study SS-31’s effect on ion leakage rates at their respective yielding *δ_TM_*, five trials were randomly selected from the unbiased simulation trials in which pore formation occurred for each lipid composition and exposure (SS-31 vs. apo). These five randomly selected trials served as the starting points for our CompEL simulations at the yielding *δ_TM_*. Specifically, structures from the randomly selected trials were output one ns before the first ion transport occurred, and each of those five states was run for seven trials (35 trials total) of 100 ns each. Above the yielding *δ_TM_*, where electroporation always occurs, we used the same initial starting point for the CompEL simulations as we used in the unbiased simulations for all 35 trials.

In sum, we generated over 400 μs of simulation, 35 μs from the CompEL simulations and 394 μs from the unbiased simulations. The unbiased and CompEL simulations were saved every 10 ps. Electrostatic and Lennard-Jones (LJ) interactions were shifted to zero at the cutoff (1.2 nm), with electrostatics shifted from 0 nm to the cutoff, and LJ interactions shifted from 1.0 nm to the cutoff. Long-range electrostatic interactions were calculated using the particle mesh Ewald method with a Fourier spacing of 0.12 nm. All double-bilayer system production runs were simulated in the NPT ensemble at 303.15 K and 1.0 bar with the Nosé-Hoover thermostat,^63,64^ Parrinello-Rahman barostat^65^ with the semi-isotropic pressure coupling scheme and periodic boundary conditions in the *X-*, *Y-*, and *Z-* directions. All hydrogen bonds were constraints using the LINCS algorithm.^66^

### Trajectory Analysis

#### Electrostatic Potential Calculations

We divided each simulation box into 500 slabs (∼0.5 Å wide) along the *Z*-dimension and calculated the average charge density in each slab using the GROMACS *gmx density* function. Charge density was calculated about the membrane’s center of mass because fluctuations of the bilayer in the *Z*-direction can lead to inaccurate estimates (i.e., smearing of the charge density).^32^ The electrostatic potential across the lipid bilayer *Ψ_z_* is related to the averaged charge density 𝜌(z) via the Poisson equation (eq. 1), where the *Z-*direction is normal to the bilayer:

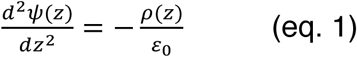

where *ε_0_* is the vacuum permittivity. The membrane electrostatic potential was calculated according to the following equation (eq. 2) ^31,32^

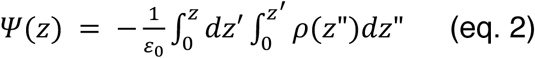

with a boundary condition *Ψ(Z=0)* = Ψ *(Z=L_z_)*, where *L_z_* is the box length in the *Z*-direction. The charge density and electrostatic potential were calculated for all atoms and also further decomposed the charge density and electrostatic potential into the averages across trials with or without electroporation. For the trials in which pores formed, we restricted our analysis to the trials from the unbiased simulations that experienced their last change in *δ_TM_* before 415 ns; unbiased simulations that experienced their last change in *δ_TM_* after 415 ns were omitted. For the qualifying trials, we used the final 50 ns of the simulation (i.e., 450-500 ns in trajectory time) allowing a minimum of 35 ns of simulation time between the last ion flux so the bilayer and ions in the interfacial region could equilibrate.

#### Electroporation and Ion/Peptide Flux Calculations

In the unbiased simulations in which pores formed, in-house Python scripts were used to compare rates of electroporation, simulation time before pore formation, and ion flux between conditions. These scripts took in atomic coordinates from the trajectory and counted the number of positively (Na^+^, SS-31: N-terminus, Arg guanidinium, Lys ammonium) and negatively charged (Cl^-^) species in each water compartment. The *Z*-component of the center of mass (Z^com^) of each bilayer was used as the upper and lower threshold between the IMS and matrix water compartments. While electroporation was visually confirmed in many trials, our metric for defining when electroporation began was a change in the *δ_TM_* relative to the initial condition. For analytical purposes, we estimated the approximate time when the pore closed — and ions re-established equilibrium in the membrane interfacial region — to be 35 ns after the last change in the *δ_TM_*. In-house Python scripts using Numpy’s polyfit function^67^ were used to perform linear and exponential regressions on the time to pore formation (τ) data. To fit an exponential curve to the data, we performed a linear regression on the transformed variables (ln(τ) vs. *δ_TM_*). To account for heteroscedasticity inherent in exponential data — where small τ values at high *δ_TM_* contribute disproportionately to the fit — we applied weights proportional to 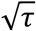. This ensured that small and large *δ_TM_* contribute more evenly to the fit, leading to a more balanced estimation of τ at smaller (i.e., more physiologically relevant) *δ_TM_* values.

The passive transmembrane transport, or ion flux, of Na^+^ and Cl^-^ was quantified by calculating the change in the number of each ion in the IMS and matrix compartments at the end of the 500 ns simulation relative to the starting conditions. Quantifying the transmembrane transport of SS-31 was slightly more involved. SS-31 is larger and often both entered and held a semi-stable position in the transient pores indicating its Z^com^ crossing the bilayer’s Z^com^ was not a reliable measure of transmembrane transport because the peptide could (and often did) return to its original compartment when the pore closed. Through an iterative process, we defined a successful transmembrane transport event as a peptide whose Z^com^ had crossed the Z^com^ of a set of lipid acyl chain atoms (POPC: C29, C39; TOCL: CA12, CB12, CC12, CD12) on the opposing leaflet and a bilayer whose pore had closed. Pore closing was defined as a bilayer which had 4 or fewer waters in the hydrophobic region of the bilayer between the above listed lipid atoms in each leaflet.

In the CompEL simulations, the CompEL module natively measures the number of Na^+^ and Cl^-^ ions which are transported from one compartment to another over time. Inhouse Python scripts were used to measure the flux rates (ion/ns). In most cases, the first 20 ns after ion flux began were used to quantify each ion’s flux rate. For trials with ≥ 2 ions fluxed of a given species in the first 20 ns after ion flux began, a linear regression was used to measure the flux rate over 20 ns (see **Figs. S5-S14** for linear regression plots for each trial). However, we adjusted the script in a small minority of trials to accurately capture some very low flux rates. For trials in which *t* ns passed between the first and second ion flux, where *t* > 20 ns, the flux rate was set to two ions per *t* ns. For trials in which one ion fluxed over the entire trajectory, the flux rate was set to 1 ion per 100 ns. Two trials with no flux from either ion were omitted from analyses of the ion flux rates in CompEL simulations.

#### Membrane Properties

We restricted our analyses of lipid headgroup angles to those trials from the unbiased simulations that did not experience electroporation to focus on how SS-31 affects these properties in stable bilayers under the influence of a *ΔΨ_m_*. The *gmx gangle* function was used to calculate the angle between a lipid headgroup vector and a unit vector in the positive (upper leaflet) or negative (lower leaflet) *Z*-direction. The lipid headgroup vector for POPC lipids is defined as the vector pointing from the phosphorus to the nitrogen atom, and for TOCL, the headgroup vector points between the two phosphorus atoms.

We included trials that did experience electroporation for bilayer thickness, and consequently membrane capacitance, to draw contrast and give insight into the capacitance after pores formed. For the trials in which pores formed, we restricted our analysis to the trials that experienced their last change in *δ_TM_* before 415 ns. For the remaining trials, we used the final 50 ns of the simulation (i.e., 450-500 ns in trajectory time) allowing a minimum of 35 ns of simulation time between the last ion flux so the bilayer could equilibrate. In-house Python scripts measuring the distance between lipid phosphates in the top and bottom membrane’s upper and lower leaflets were used to estimate the bilayer thickness. The membrane specific capacitance (*C_s_*) was estimated by calculating the specific capacitance of the bilayer’s hydrophobic region (*C_h_*) according to eq. 3^68^

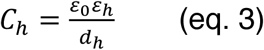

where *d_h_* is the width of the bilayer hydrophobic region and 𝛆*_h_* the dielectric constant of the hydrophobic region. We used the distance between lipid phosphates for *d_h_* and 𝛆*_h_*=2.61, as previously reported.^68^ The total membrane capacitance (*C_m_*) was then calculated by multiplying *C_s_* by the surface area of the bilayer plane, which we quantified as the *X-Y* box dimensions.

#### Transverse Atom Density

We measured the transverse atom density of unbiased ion imbalance simulation trials which did not experience electroporation. We divided each simulation box into 500 slabs (∼0.5 Å wide) along the *Z*-dimension and used the GROMACS *gmx density* function to calculate the average mass density of various subgroups of atoms. Atom densities were calculated about and symmetrized around the *Z*-coordinate of the central water compartment’s (i.e., the “IMS”) center of mass (COM). The Na^+^ and Cl^-^ densities were normalized to the average bulk density at the edge of the box. The normalized water hydrogen: oxygen (H:O) density difference was calculated by subtracting the normalized densities 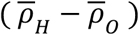, where the normalized densities 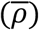 are calculated by dividing the mass density by each element’s atomic mass and then dividing the hydrogen density by two to create a metric in which a value of 0 represents a randomly oriented (non-polarized) layer of water.

#### Data Analysis and Presentation

Unless stated otherwise, individual trials were treated as independent samples. Individual lipids were treated as independent samples for the lipid angle analyses. All plots showing means use the means ± the standard deviation or standard error depending on whether we are trying to illustrate the width of the distribution or uncertainty in the measurement. Trajectories were processed for Python analysis using MDTraj.^69^ The contingency chisquare test was used to test for significance in the proportions of electroporation from different conditions. All images of the systems were created using VMD.^70^ The Matplotlib Python package was used to create various plots and figures.^71^

We used multivariate linear modeling to examine how ion fluxes were influenced by the experimental conditions, performed using R software run in R Studio.^72^. We tested whether the combination of transmembrane potential (𝝳_TM_), peptide, and lipid composition had significant effects on the fluxes of chloride (Cl⁻) and sodium (Na⁺) ions by fitting a multivariate linear model to the square roots of the Cl⁻ and Na⁺ flux values, using all main effects and interactions among 𝝳_TM_, peptide, and lipid as predictors. The square root transformation was applied to stabilize variance across conditions. This approach allowed us to assess whether any of the experimental factors had a statistically significant effect on the combined ion flux profile, rather than analyzing each ion separately. We were also able to use the model to examine effects of specific parameters on estimated marginal means. The multivariate model in R was analyzed using the Anova function from the CAR package (type III SS).^73^ The package emmeans was used to analyze marginal effects and to generate interaction plots.^74,75^

## Results

### Peptides and CL increase Cm and reduce *ΔΨm*

We have employed all-atom double bilayer MD simulations with four increasingly large *δ_TM_* values to determine how the presence of CL and SS-31 influence bilayer properties, membrane electroporation, and ion flux under an imposed transmembrane electrochemical imbalance. Our initial analyses focus on systems in which electroporation does not occur, to confine our measurements to systems with intact bilayers. From this data set, we observe that the presence of CL slightly increases bilayer thickness (**Fig. 2A**), increases bilayer area (**Fig. 2B**) and increases total membrane capacitance (*C_m,_* **Fig. 2C**). Addition of CL has been shown to have similar effects on bilayer properties to previous MD studies,^12,76^ though we’re unaware of any studies examining the effect of CL under explicitly-modeled *ΔΨ_m_* potentials. We also observe a decrease in thickness, increase in area, and increase in *C_m_* corresponding with increasing 𝝳_TM_ across both lipid compositions.

**Figure 2.**
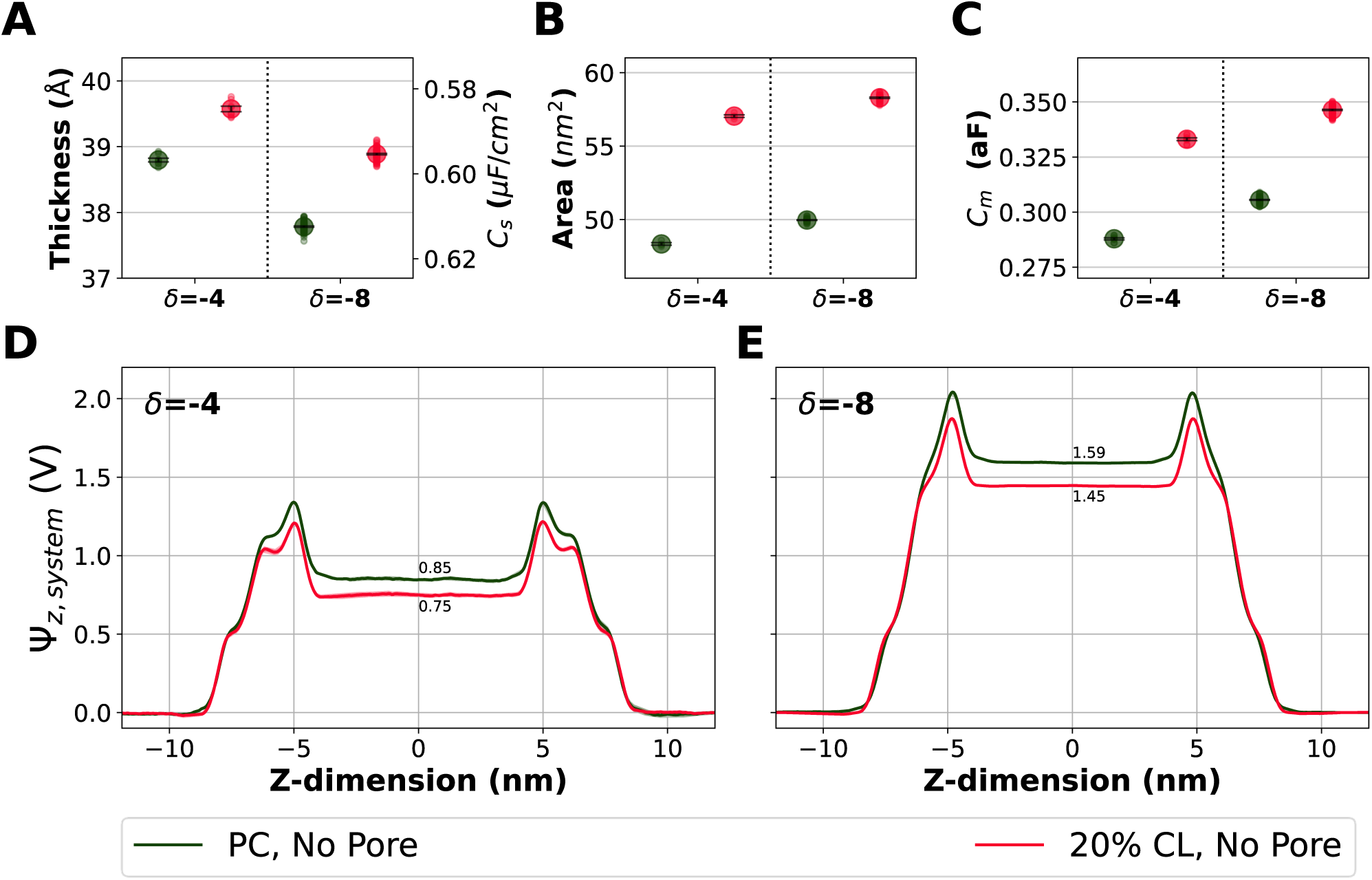
Electrostatic potential and membrane properties for intact bilayers with and without cardiolipin. Comparison of the average bilayer thickness (A, *left axis*), specific capacitance (A, *right axis*), bilayer surface area (B), and total membrane capacitance for the all POPC and 20% CL bilayers (without peptides present) (C). The small, color-coded circles represent the averages from each bilayer in individual simulations (see Fig. 1D *n* in each condition), and the larger circles with errors represents the means ± the standard errors. Comparison of electrostatic potential between the all PC and 20% CL bilayers at *δ_TM_* =-4 (D) and *δ_TM_* =-8 (E). The dark lines represent the average across *n* trials and the lighter shaded regions represent averages ± the standard error.

In CL-containing bilayers, *δ_TM_* values of −4, −8, and −12 respectively yielded transmembrane potential (*ΔΨ_m_* = *Ψ_z_*(*Z*=0)) values of 0.75 V, 1.45 V, and 1.98 V. Meanwhile in all POPC bilayers, the same *δ_TM_* values of −4 and −8 yield larger *ΔΨ_m_* values of 0.85 V and 1.59 V (**Fig. 2D-E**). The observation that CL containing bilayers exhibited *ΔΨ_m_* values 100-150 mV lower than pure PC bilayers, for the same *δ_TM_* can be explained by the higher capacitance of CL containing bilayers. Given the molecular properties of CL, a bilayer with 20% 4-tailed lipids should naturally have more surface area than one with 100% 2-tailed lipids, and thus greater total capacitance. Further, our results show that increasing *δ_TM_* correlates with increased *C_m_*, a result which is probably due to electrocompression (i.e., reduction of distance between head groups driven by the electrostatic attraction of ions across the bilayer).^77^ This leads to lateral expansion of the membrane and a decrease in the non-polar thickness, hence increased capacitance. This can be seen by examining the changes in membrane area (increasing), thickness (decreasing), and *C_m_* (increasing) in systems which did not porate (**Fig. 2, Fig. S15**). These results underline how electrochemical stress causes bilayers to elastically deform their dimensions — increasing *C_m_* but also inducing area strain — until electroporation becomes more energetically favorable than further elastic deformation.

Adding SS-31 to the double-bilayer systems had a similar effect to the presence of CL. Systems containing SS-31 exhibited *ΔΨ_m_* values 50-100 mV smaller than the peptide-free (*Apo*) controls (**Fig. 3B, S16**). Membrane binding of SS-31 increases a given bilayer’s surface area and decreases its bilayer thickness (**Fig. S15** *top row*), as we observed in our previous studies.^12–14^ Accordingly, SS-31’s presence results in a ∼1-3% increase in the membrane specific capacitance (**Fig. 3A**). With an increase in *C_m_* and constant *δ_TM_*, the theoretical prediction is that SS-31 causes a decrease in *ΔΨ_m_*, which is what is directly observed in the simulations at *δ_TM_* values of −8 and −12 (**Fig. 3B**). This increase in capacitance is consistent with our previous computational study^14^ and supported by our previous experimental data suggesting SS-31 can prevent hyperpolarization of the IMM.^12^ These results lend support to the hypothesis that SS-31 increases ATP production by increasing the amount of potential energy which can be stored across the IMM without hyperpolarizing.

**Figure 3.**
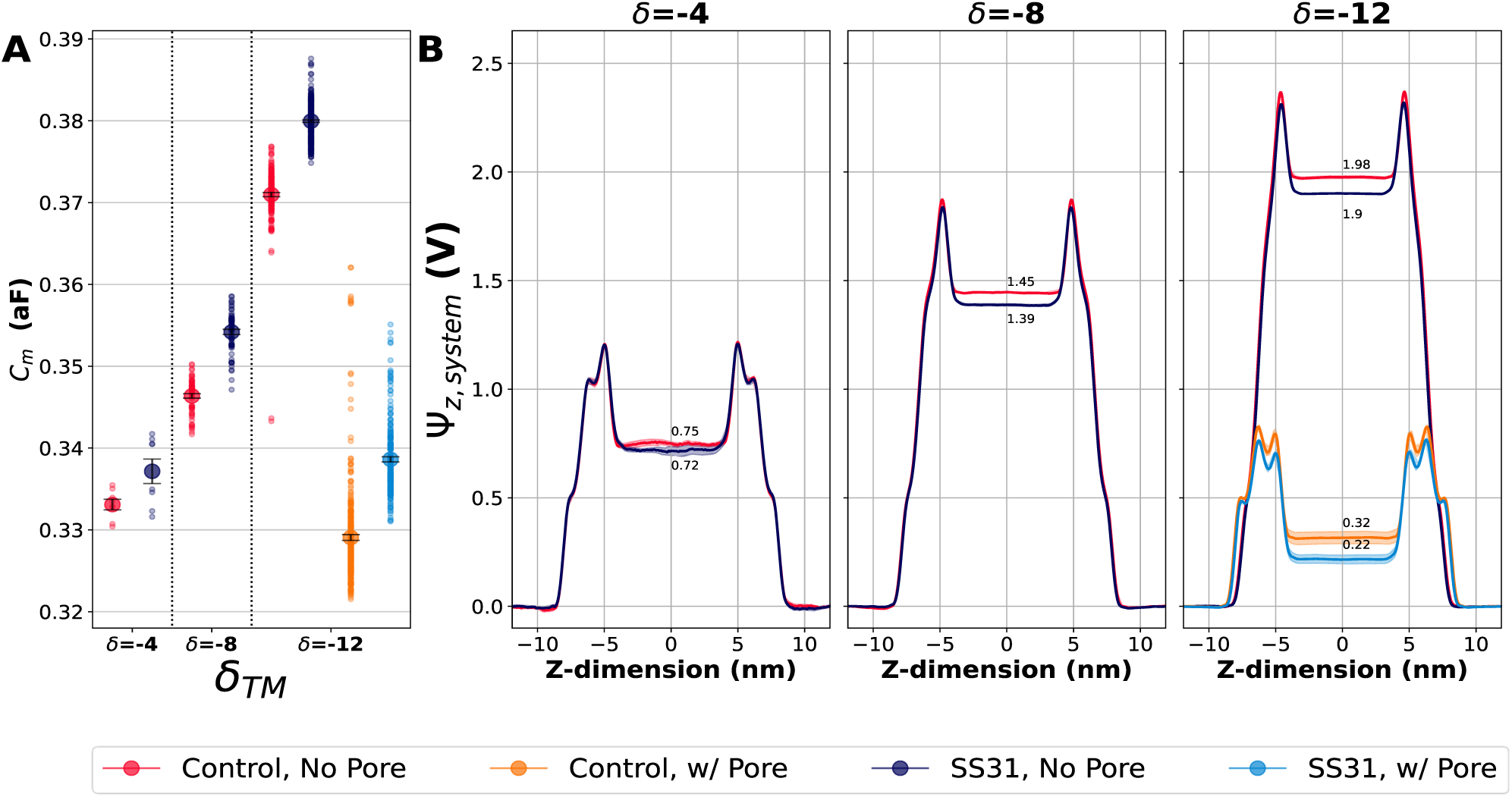
Electrostatic potential and membrane properties in CL-containing bilayers with and without SS-31. (A) Comparison of the average total membrane capacitance (*C_m_*) for CL-containing bilayer conditions. The small, color-coded circles represent the averages from each bilayer in individual simulations (see Fig. 1**D**’s 20% CL subsection for *n* in each condition), and the larger circles with errors represents the means ± the standard errors. (B) Comparison of electrostatic potential (Ѱ_z_) between the apo bilayers with 20% CL and the SS-31-containing systems at different levels of exposure, as well as with and without passive ion leakage. The dark lines represent the averages across *n* trials and the lighter shaded region represent averages ± the standard error.

### SS-31 reduces electroporation propensity in CL-containing bilayers

We next investigated systems in which electroporation does occur to quantify the effects of addition of CL and/or SS-31. Trials with electroporation were identified by detecting a change in the *δ_TM_* compared to the initial condition. As expected, electroporation was not observed at the timescales of our simulations at the lowest *δ_TM_* (−4, **Table 1**, **Fig. 4A**), regardless of lipid composition or peptide exposure. This is likely because electroporation is a rare event at *ΔΨ_m_* voltages below the threshold range.^29^ In the peptide-free bilayer systems, the addition of CL to shifted the *δ_TM_* required for electroporation, indicating CL is protective against a large *ΔΨ_m_*. At a *δ_TM_* of −8, electroporation was similarly rare for both lipid compositions, where bilayers did not porate in CL containing bilayers and occurring only once in the all-POPC bilayers. At the higher *δ_TM_* strength of −12, CL-containing bilayers electroporated in 188/286 trials (66%) significantly less frequently (p = 0.0104, Fisher’s exact test) compared to the all-POPC bilayers which porated in all trials (12/12).

**Figure 4.**
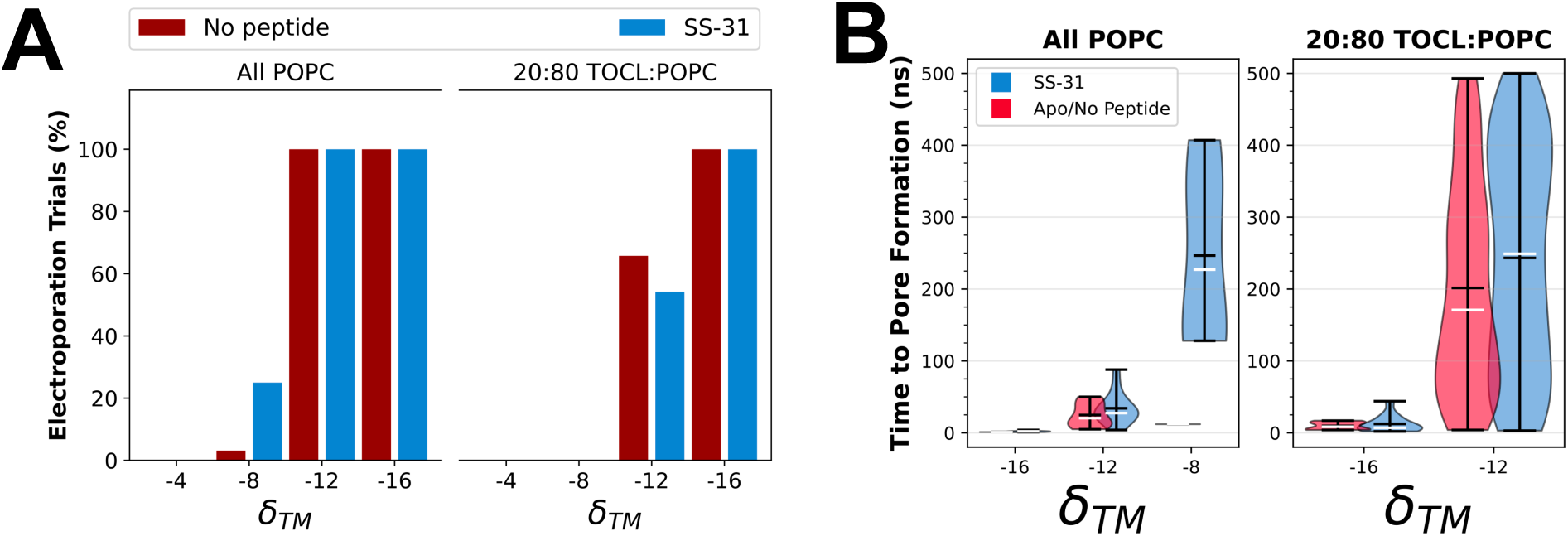
Comparison of electroporation propensities and kinetics between conditions. (A) Comparison of the percent of simulations in which ion leakage occurred between bilayer lipid compositions, applied voltage (*δ_TM_*), and the presence or absence of SS-31. The contingency chi square and fisher’s exact tests were used to test for significance (✱ indicates p < 0.05). (B) Comparison of the time lapse before pore formation between different *δ_TM_* levels in all PC (*left*) and 20% CL (*right*) bilayer systems with (*blue*) and without (*red*) SS-31. Black horizontal lines in the center of the error bars represent the average, white horizontal lines represent the median, and black lines at the end of the error bars represent the largest or smallest individual values. Shaded violin areas represent the distribution of values for each condition. Focused comparison of the amount of time before pore formation for the all PC bilayer systems. Data was pooled from the simulations with and without SS-31. Large black dots represent the average time to pore formation for a given *δ_TM_* while the smaller black dots represent individual trials. Error bars are the standard deviation. The raw data were fit to both a linear and exponential curve to predict how the relationship between *δ_TM_* and time to pore formation may behave at lower *δ_TM_* values.

**Table 1.**
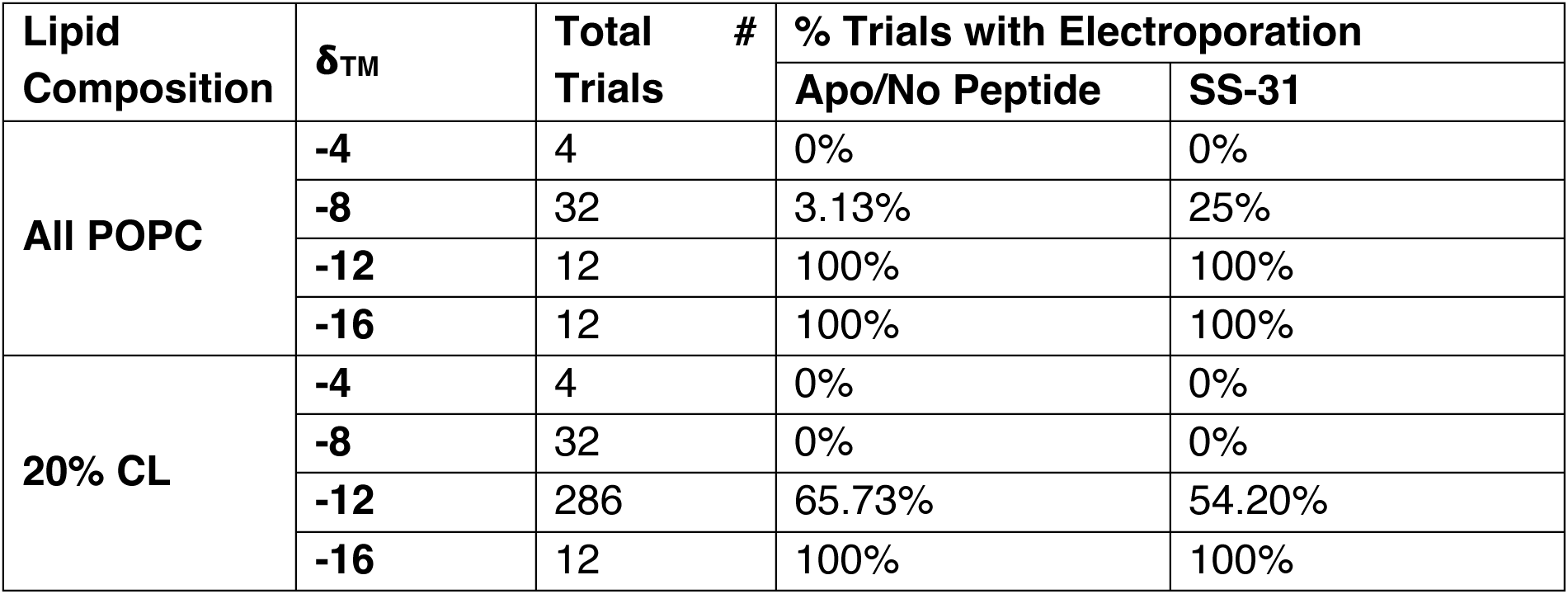
Comparison of electroporation rates between PC vs. 20% CL and Apo vs. SS-31 conditions. Each trial represents a 500 ns unbiased (non-CompEL) MD simulation.

In the presence of SS-31, at a *δ_TM_* of −8, electroporation occurred in 25% (8*/* 32) of trials for the all-POPC bilayer but none of the trials for the 20% CL bilayer (0/32, p =0.0047). At the higher *δ_TM_* strength of −12, all POPC bilayers experienced electroporation in all trials (12/12) while the 20% CL bilayer electroporated in 54% of trials (155/ 286, (p = 0.0015, Fisher’s exact test). Both lipid compositions electroporated in all trials at the highest 𝝳_TM_ strength of −16. These results are consistent with Lin & Alexander-Katz’s finding that bilayers electroporate sporadically in the “yielding” *δ_TM_* but always at or above the “threshold” *δ_TM_*. However, our results differ in the precise quantitative relationship between charge imbalance and electroporation. The *δ_TM_* values of −4, −8, −12, and −16 would respectively equate to *e*/Lip values (and regimes of) of 0.053 (linear), 0.107 (threshold), 0.160 (threshold), and 0.213 (threshold). Based on these *e*/Lip values and a strict interpretation of Lin-Alexander-Katz’ three regimes, the prediction would be that electroporation occurs within 80 ns for all systems at or above a *δ_TM_* of −8. This is inconsistent with our results; the PC and 20% CL bilayers exhibited different electroporation propensities and kinetics, suggesting that the linear, yielding, and threshold regimes for electroporation may be unique to a given bilayer’s lipid and protein composition. This different behavior between the PC and CL-containing bilayers is why we chose to use *δ_TM_* over *e*/Lip; *e*/Lip is a compelling metric because it captures the relationship between electroporation and membrane surface area, but it seems inappropriate for comparing membranes with differing lipid and/or protein compositions.

To better understand how *δ_TM_*, lipid composition, and peptide exposure influence pore formation kinetics in all-atom MD simulations, we measured the elapsed time before pore formation (τ) under various conditions (**Fig. 4B**). Our results indicate that the magnitude of *δ_TM_* is inversely correlated with τ, consistent with previous findings by Lin & Alexander-Katz on coarse-grained lipid bilayers. Further, the CL-containing bilayers at *δ_TM_* =-12 had similar average *τ* compared to the all PC bilayers at *δ_TM_* =-8, and the addition of SS-31 yielded a modest increase in average τ compared to the peptide-free bilayers. These results are consistent with effects CL and SS-31 have on *ΔΨ_m_* and *C_m_*.

### SS-31 effects are sensitive to bilayer composition

SS-31 increases the *C_m_* in both lipid compositions (**Fig. 3A**), and the expectation is that it would decrease electroporation propensity in both bilayer compositions as well. However, we found the effect of SS-31 on electroporation to be affected by lipid composition. At a *δ_TM_* of −12 in 20% CL bilayers, electroporation occurred significantly (p=0.006) less often in the presence of SS-31 (155/286 trials, 54.2%) than in the peptide-free control (188/286 trials, 65.7%) (**Fig. 4A**, **Table 1**). However, at a *δ_TM_* of −8, the yielding *δ_TM_* range for PC systems, electroporation occurred significantly (p=0.031) more often in the presence of SS-31 (8/32 trials, 25%) than in the Apo control (1/32 trials, 3.1%). There are several possible explanations for this behavior which are presented in the **Discussion** section.

### Peptides and lipid composition alter passive transmembrane ion flux rates

After observing that both lipid composition and peptides alter the probability of electroporation, we used computational electrophysiology (CompEL) simulations^36,58,62^ — to maintains a constant *ΔΨ_m_*, analogous to a voltage-clamp experiment — to investigate whether CL and/or SS-31 can influence the rates of cation or anion transport across the bilayer. CompEL simulations have two main advantages for measuring ion flux over the unbiased simulations. First, by using high *δ_TM_* strengths and careful selection of initial states, we were able to guarantee electroporation would occur on the simulation timescale, thus allowing the ability to measure CL’s/SS-31’s direct effect on ion flux independent of pore formation propensity. Second, since CompEL simulations maintain a constant *δ_TM_*, ions leak across the bilayer in greater frequency and quantity facilitating the measurement of flux rates and more robust statistical analysis. The raw rates of ion flux for each condition are displayed in **Table 3** and **Figure S17.**

**Table 3.** Rates of ion flux from CompEL simulations. The CompEL simulations were run for 35 trials. In the all POPC system with SS-31 at a *δ_TM_* −8, two trials were omitted from the below table because no flux of either ion occurred during the 100 ns simulation.

| **Lipid Composition** | **$\delta_{TM}$** | **Na<sup>+</sup> Flux (ion/ns ± SE)** |  | **Cl<sup>-</sup> Flux (ion/ns ± SE)** |  |
| --- | --- | --- | --- | --- | --- |
|  |  | **SS-31** | **No peptide** | **SS-31** | **No peptide** |
| **20% CL** | **-12** | 0.64 ± 0.05 | 1.58 ± 0.12 | 0.08 ± 0.02 | 0.16 ± 0.03 |
| | **-16** | 1.66 ± 0.17 | 3.15 ± 0.22 | 0.43 ± 0.08 | 0.85 ± 0.1 |
| **All POPC** | **-8** | 0.14 ± 0.02 | 0.3 ± 0.03 | 0.22 ± 0.03 | 0.29 ± 0.03 |
| | **-12** | 0.24 ± 0.04 | 1.03 ± 0.04 | 1.2 ± 0.11 | 1.84 ± 0.08 |
| | **-16** | 0.31 ± 0.05 | 2.29 ± 0.05 | 2.27 ± 0.15 | 4.36 ± 0.08 |

To deepen our understanding of how ion flux varies across conditions, a multivariate ANOVA (MANOVA) model describing Na^+^ and Cl^-^ fluxes as a function of *δ_TM_*, lipid composition, peptide exposure, and statistical interactions between them was used to analyze the data. MANOVA was used because Na^+^ and Cl^-^ fluxes are paired for each instance of pore formation. Initial linear models showed considerable heteroskedasticity; to reduce it, a square root transform was applied to the independent variables (**Fig. 5A**). This model fit the data well (**Fig. 5B**) and accounted for ∼80% of total variance (adjusted r^2^ for Cl^-^ flux = 0.84, adjusted r^2^ for Na^+^ flux = 0.78). Further analysis of separate ANOVA tables for Na^+^ and Cl^-^ fluxes indicated that there were several significant interaction effects, including a significant three-way interaction between *δ_TM_*, lipid, and peptide for Cl^-^ flux, which complicated interpretation of main effects. Analysis of marginal effects at the average *δ_TM_* (−12.8) supports several conclusions detailed below.

**Figure 5.**
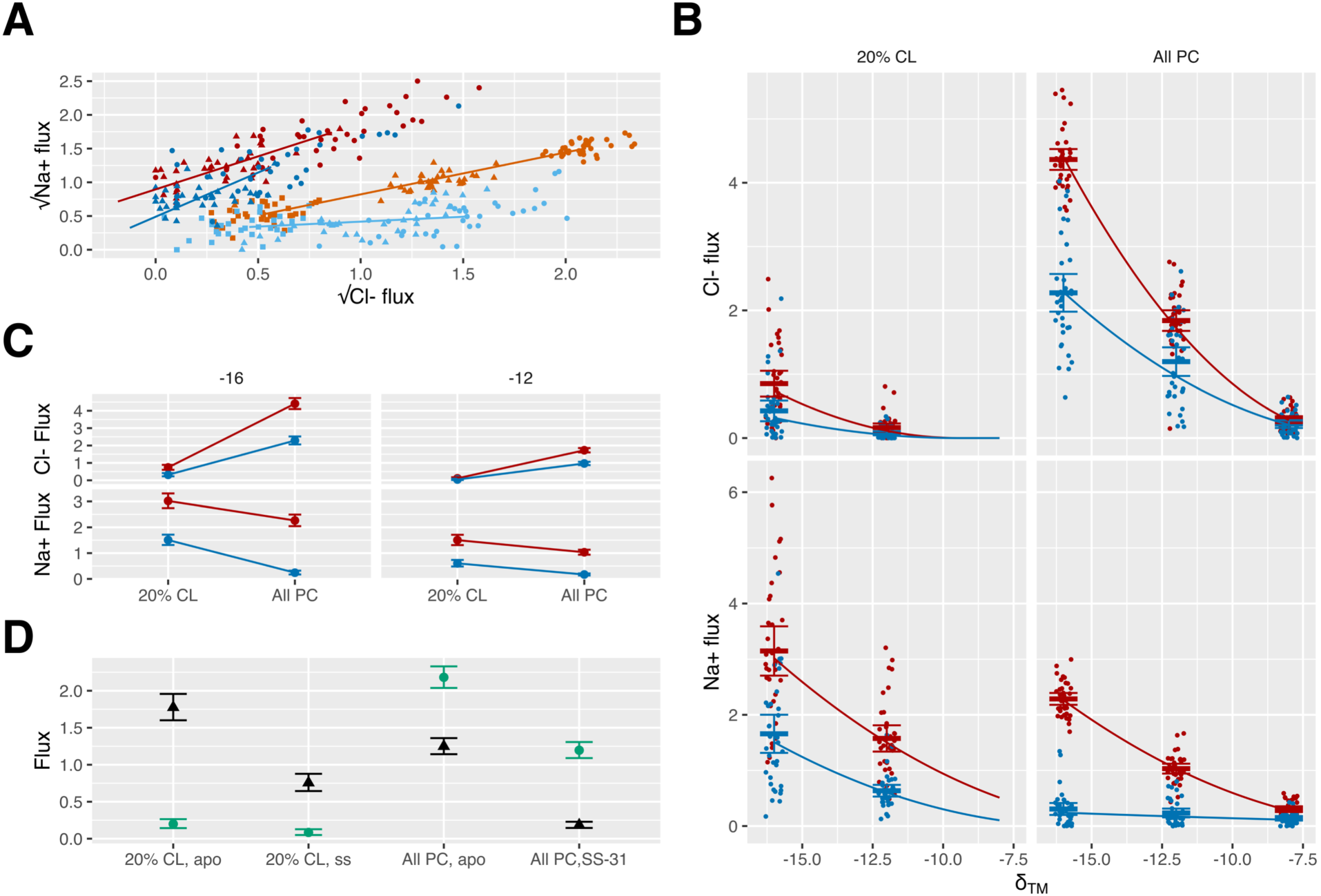
Multivariate analysis of CompEL ion imbalance simulations. A) Plots of the square root of Na^+^ flux vs. the square root of Cl^-^ flux in the apo control condition (red traces) or in the presence of SS-31 peptide (*blue traces*) in all-POPC membranes (*dark blue*, *bright red*) or membranes containing 20% CL (light blue, purple red) at 𝝳_TM_ of −8 (*squares*), −12 (*triangles*) or −16 (*circles*). Lines are fits to the model. B). Plots of fluxes on the linear scale vs 𝝳_TM_ in the apo control condition (*red*) or in the presence of SS-31 peptide (*blue)*. Lines are fits to the model. C) Plots at *δ_TM_* of −12 (*left*) and −16 (*right*) illustrating key interactions in the model. D) Estimated marginal values calculated at −12.8 *δ_TM_* for Cl^-^ (green *circles*) and Na^+^ (black *triangles*) flux. Only two of the possible pairwise comparisons (1: 20% CL apo Cl^-^ vs all PC SS-31 Na^+^, and; 2: all PC Apo, Na^+^ vs All PC, SS-31, Cl^-^) are not significant at p < 0.05 or smaller after correction for multiple testing.

First, Na^+^ flux is significantly higher in membranes containing 20% cardiolipin than zwitterionic all POPC membranes (**Fig. 5B, bottom, Fig. 5D**), while the opposite is true for Cl^-^ flux (**Fig. 5B, top, Fig. 5D**). While 20% CL bilayer systems have more Na^+^ ions than the analogous all POPC bilayer systems, to balance CL negative charge (**Figs. S1-S4**), a mechanistic explanation is also plausible. Cardiolipin imparts a negative charge onto the bilayer surface, drawing Na^+^ ions closer to the bilayer (**Fig. 6A** *top graphs*). When a pore forms, Na^+^ ions, being more proximal to the pore, may be more likely to transport than more distal Cl^-^ ions. The inverse can be said for the all POPC system which exhibits stronger accumulation of Cl^-^ ions near the bilayer (**Fig. 6A** *middle graphs*).

**Figure 6.**
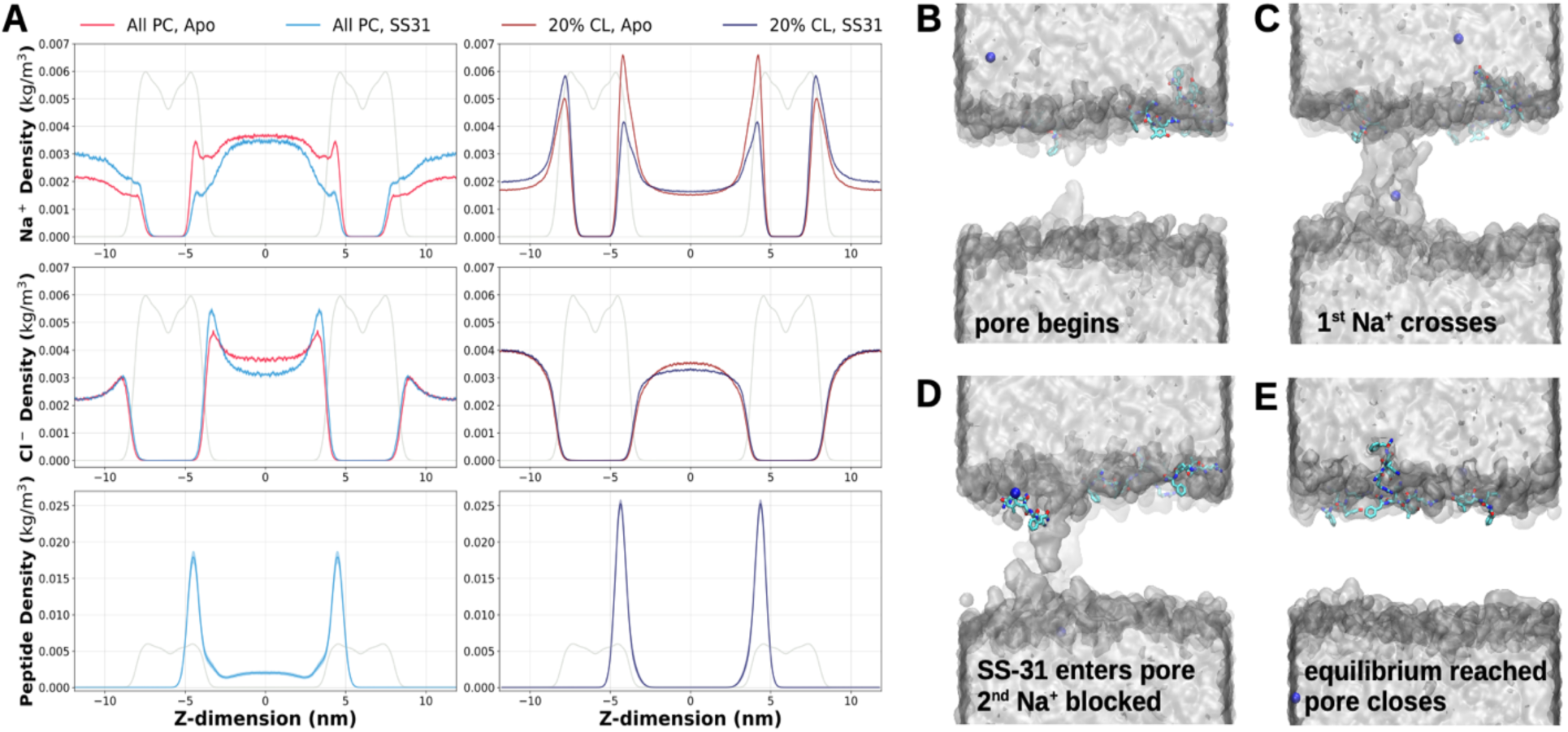
Transverse atom density profiles and SS-31’s effect on ion distributions. (A) Transverse atom density profiles of sodium ions (top), chloride ions (middle), and peptides (bottom) for the all POPC (left) and 20% CL bilayers (right) along the Z-dimension (i.e., normal to the membrane) in relation to the system’s COM (i.e., Z=0 represents center of IMS compartment). The light gray line in all graphs represents the transverse atom density of the bilayers. Note that the transverse atom density profiles were symmetrized about Z=0, which corresponds to the midpoint between the two membranes.. (B-E) Four snapshots of a system in which ionic imbalance (Na^+^ = *blue spheres*) leads to the spontaneous formation of a solvated (silver surface) nano-pore in the bilayer (lipids = hidden to improve pore visualization) where a peptide (wireframe: carbon = *cyan*, nitrogen = *blue*, oxygen = *red*) partially occluded the pore impeding Na^+^ flux.

Second, SS-31 significantly decreased both Na^+^ and Cl^-^ ion flux compared to the no peptide control systems (**Fig. 5A, 5B, 5D**). This is consistent with SS-31’s observed ability to increase *C_m_* and reduce *ΔΨ_m_*, which would decrease the drive for spontaneous ion flux. Third, SS-31 inhibited Na^+^ flux about twice as much as Cl^-^ flux, but the effects were complicated by interactions between lipid composition and *δ_TM_* (**Fig. 5C**). When *δ_TM_* was −16, both Na^+^ and Cl^-^ fluxes are reduced more by SS-31 in all POPC membranes than in membranes containing 20% cardiolipin. When *δ_TM_* was −12, Cl^-^ flux was still reduced more in all POPC membranes than in membranes containing 20% CL, but the reduction in Na^+^ flux was equivalent.

SS-31’s depletion of Na^+^ ions and enrichment of Cl^-^ ions in the membrane interfacial region (**Fig. 6A**) may explain why the peptide had a larger impact on reducing Na^+^ flux than Cl^-^ flux. Notably, SS-31’s effect on ion flux extended to the all POPC systems despite the peptide’s promotion of electroporation, suggesting that SS-31’s ability to slow ion flux may be independent from its effect on electroporation in all POPC bilayers.

### Pore formation precedes passive transmembrane transport of SS-31

When visualizing trajectories from unbiased simulations in which pores had formed, we observed something somewhat unexpected in many trials, transmembrane transport of SS-31. MTPs are known to accumulate at or near the IMM of mitochondria across a variety of cell types,^15,16^ meaning they must cross both the cell membrane and OMM but there is little consensus about the mechanism of how MTPs’ transport across bilayers. Previous studies on potential transmembrane transport mechanisms for cell-penetrating peptides (CPPs), which have similarly cationic and aromatic sequence compositions, support mechanisms involving the formation of aqueous pores.^78^ However, there has been limited direct evidence on the mechanism MTPs use to cross lipid bilayers. Understanding the mechanism of membrane transport is of interest to basic science and may be important for future drug optimization. A recent study employed steered MD simulations to characterize the energetics of transporting a bio-conjugated MTP through a lipid bilayer and reported a very high energy barrier, suggesting that SS-31’s spontaneous transport across the bilayer may be a rare event.^79^ This is consistent with our previous published work on SS-31.^12–14^ Unassisted transmembrane transport of SS-31 did not occur in any of our previous studies in which we used single bilayer simulations, nor did it occur in this study at *δ_TM_* values below the apparent yielding range for electroporation for each lipid composition (**Table 4**).

**Table 4.** Fraction of trials from unbiased ion imbalance simulations with transmembrane transport of SS-31. Note that electroporation did not occur in any 20% CL bilayer at 𝝳_TM_=-8 which is why they are listed as N/A.

| Lipid Com-<br>position | Trial<br>Selection | Number Trials w/ Transmembrane Transport of SS-31 |  |  |
| --- | --- | --- | --- | --- |
| | | $\delta_{TM}=-8$ | $\delta_{TM}=-12$ | $\delta_{TM}=-16$ |
| All POPC | w/ pores | 2/8 (25.0%) | 4/12 (33.3%) | 6/12 (50.0%) |
|  | All | 2/32 (6.3%) | 4/12 (33.3%) | 6/12 (50.0%) |
| 20% CL | w/ pores | N/A | 25/155 (16.1%) | 4/12 (33.3%) |
|  | All | N/A | 25/286 (8.7%) | 4/12 (33.3%) |

We observed passive transmembrane transport of SS-31 in all systems at or above the yielding *δ_TM_* for electroporation, though with varying frequency. Transport events were more common in the all-POPC bilayers than in the 20% CL bilayers, possibly due to favorable electrostatic interactions between SS-31 and CL headgroups,^14^ or the larger pool of unbound peptides in the POPC systems (**Fig. 6A**). Interestingly, in the standard double-bilayer simulations, only a single peptide was ever observed translocating in a given trial, after which the pore would typically close. This suggests that SS-31 transport reduces *δ_TM_* enough to destabilize the pore and prevent further translocation. However, this may be a finite-size effect: in the CompEL simulations, which maintain a constant *δ_TM_*, multiple peptide transport events were frequently observed. These observations suggest that membrane poration and peptide translocation dynamics can be strongly influenced by system size and charge distribution method.

A representative pathway for SS-31’s transport is presented in **Fig. 7**. The process starts with a peptide proximal (**Fig. 7A**) to the nucleation point for a solvated pore (**Fig. 7B**) followed by SS-31 being drawn into the pore by its lysine residue interacting with lipid phosphates from the opposing leaflet (**Fig. 7C**). From there the peptide adopts a quasistable intra-pore position with basic residues in the solvent and aromatics in the acyl chain region (**Fig. 7D**), the Arg residue then flips down to interact with lipid phosphates in the opposing leaflet (**Fig. 7E**). The peptide maintains interactions with the opposing leaflet while lipid phosphates begin to recede from the bilayer’s hydrophobic core (**Fig. 7F**) and ultimately the pore closing occurs with SS-31 stably bound on the other side of the membrane (**Fig. 7G**). To the best of our knowledge, these unbiased ion imbalance simulations represent the first computational or experimental evidence of SS-31 spontaneously transporting across a model membrane (albeit under an elevated *ΔΨ_m_* above physiological conditions). This suggests that if SS-31 does transport across bilayers without the assistance of a protein translocase, fluctuations in local *ΔΨ_m_* followed by transient electroporation may be a necessary precursor for transport, as SS-31 does not induce pores on its own.

**Figure 7.**
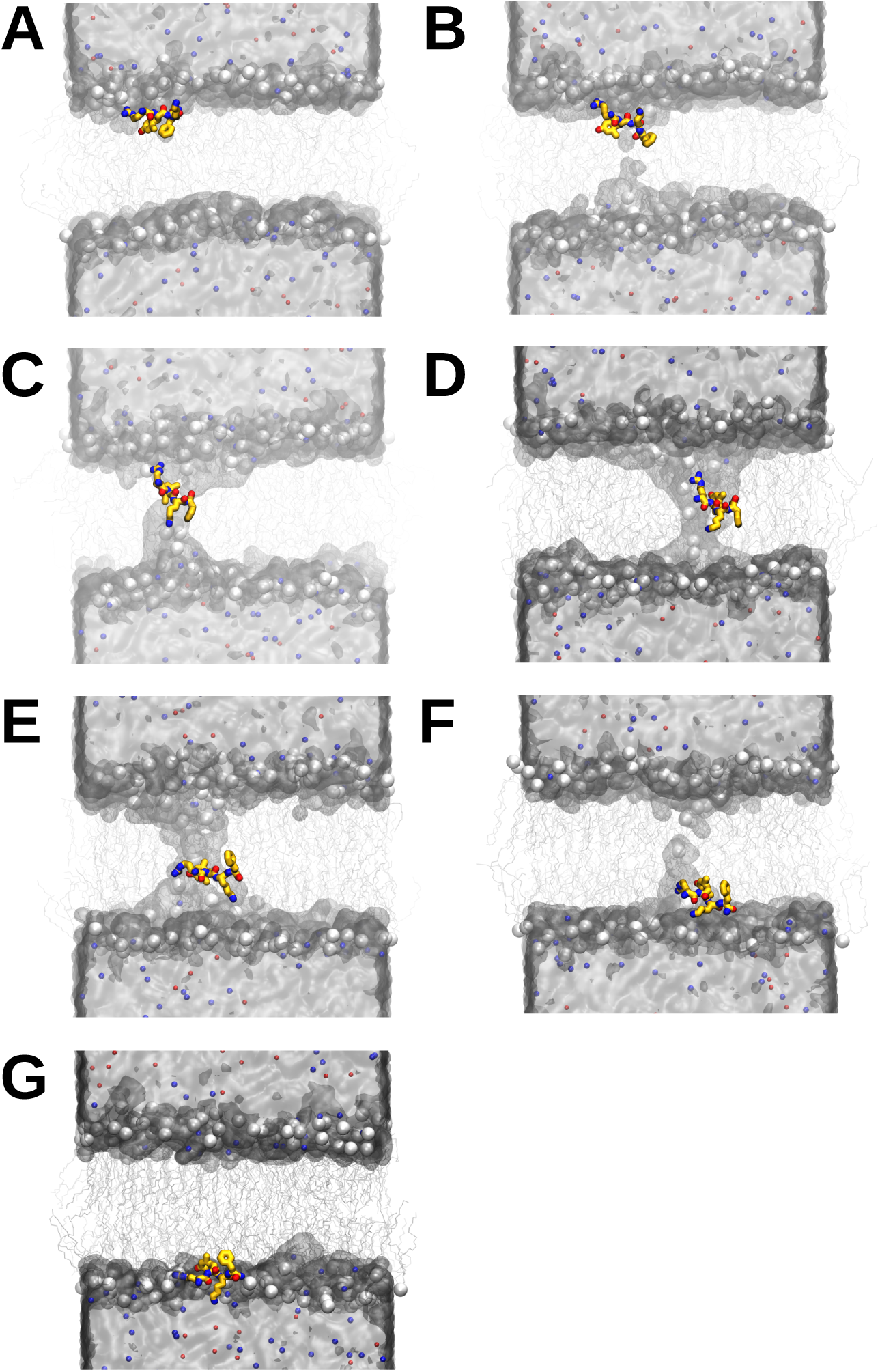
Representative pathway of passive transmembrane transport of SS-31 from unbiased ion imbalance MD simulations. (A-G) Representation of a system in which ionic imbalance (Na^+^ = blue spheres; Cl^-^ red spheres) lead to the spontaneous formation of a solvated nano-pore in the bilayer (lipid phosphates = white spheres; lipid acyl chains = gray wireframe) followed by the complete passive transmembrane transport of a peptide (wireframe: carbon = *goldenrod*, nitrogen = blue, oxygen = red) from the IMS water (silver surface) compartment to the matrix compartment.

## Discussion

### Why is CL protective against electroporation?

CL appears to stabilize bilayers against electroporation, as higher *δ_TM_* values are required to induce pore formation in the 20% CL systems. Several factors may contribute to this protective effect. First, bilayers containing CL exhibit an estimated membrane capacitance (*C_m_*) that is ∼10–15% higher than that of pure POPC bilayers. This increased capacitance reduces the transmembrane potential (*ΔΨ_m_*) generated at a given charge imbalance, thereby decreasing the electrostatic drive for electroporation. Second, surface charge density differs between the systems. POPC is zwitterionic and electrically neutral, whereas CL carries a net charge of −2. Favorable electrostatic interactions between CL phosphate groups and PC choline headgroups may raise the energetic barrier to pore formation.^80–82^ Indeed, lipid headgroup interactions have been hypothesized to influence electroporation kinetics.^83^ Third, differences in lipid geometry may also play a role. CL has a negative spontaneous curvature due to its inverted conical shape, which may resist the formation of toroidal pores — structures that require local positive curvature (as seen in **Fig. 7C–E**). ^84^ This may explain why bacteria enriched in CL show increased resistance to membrane-disrupting antimicrobial peptides.^85^ Paradoxically, however, CL’s geometry may also promote electroporation under certain conditions. Its ability to induce transient lipid packing defects^86^ can expose the membrane’s hydrophobic core to solvent, providing nucleation sites for pore formation. ^87^ Overall, the increased *C_m_* in CL-containing bilayers offers the most direct explanation for their resistance to electroporation, though contributions from lipid charge and curvature may also be contributing factors.

### Why does SS-31 affect CL bilayers differently than pure PC bilayers?

SS-31 promoted electroporation in all-POPC membranes but inhibited it in bilayers containing 20% CL. One potential explanation involves limitations in how we estimate membrane capacitance (*C_m_*), which assumes a parallel-plate capacitor model. Unlike ideal capacitors, biological membranes are dynamic and electrostatically heterogeneous. Both surface roughness^88^ and lipid composition^89,90^ have been shown to affect capacitance experimentally, but these factors are not explicitly included in our estimated *C_m_*. Moreover, electroporation is not analogous to current flow through a connected circuit — it is more akin to rupturing a capacitor plate, allowing the potential to dissipate. This highlights the importance of bilayer mechanical stability in determining electroporation propensity.

Accordingly, SS-31’s reduction of electroporation propensity in the 20% CL system could partially be attributed to its unique effects on CL-containing bilayer structural properties and membrane defects. Our previous work showed that SS-31 can occlude lipid packing defects and shield the acyl chain region of CL-rich membranes from solvent exposure,^12,13^ potentially lowering the number of transient nucleation sites that facilitate pore formation. Other potential explanations could include the different electrostatic environments of the membrane interfacial region between a bilayer with and without CL. SS-31 concentrates more strongly at the surface than Na^+^ ions which may create considerable charge repulsion in all POPC bilayer, potentially destabilizing membrane structural integrity.

The differential effects of SS-31 may also reflect limitations in our MD simulation setup.. First, SS-31’s interaction with pure POPC liposomes was just barely detectable in our experimental work using isothermal titration calorimetry.^12^ Yet in these MD simulations, SS-31 readily binds POPC bilayers, though more weakly than the CL-containing bilayers (see the decreased peaks at the bilayer surface (Z=±5) and increased atom density in the solvent (Z=-3 to Z=3), **Fig. 6A *bottom row***). This could be due to a greater peptide concentration in our simulations compared to the experimental conditions or limitations in the force field parameters, which may overestimate SS-31 binding POPC bilayers. While this reduces direct comparability to experimental conditions, POPC remains a valid control for assessing compositional effects on electroporation.. Second, finite-size effects in our bilayer simulations may influence peptide–membrane interactions and poration propensity. When SS-31 binds to IMS-facing leaflets, both inner and outer leaflets expand due to semi-isotropic pressure coupling, leading to area strain in the matrix-facing leaflet. Though such strain could occur in real membranes, its extent may be exaggerated in the finite simulation box. Notably, this effect occurs in both bilayer systems. In the 20% CL membranes, SS-31’s combined impact on *C_m_* and defect suppression appears to protect against poration. In contrast, in the all-POPC membranes, any beneficial *C_m_* effects may be counteracted by SS-31-induced destabilization of bilayer structure.

### Physiological Implications

Our results offer insight into one of SS-31’s known effects on mitochondria, its ability to reduce proton leak,^91^ an energy sink for ATP production. SS-31’s depletion of cations in the membrane interfacial region is functionally consistent with SS-31’s ability to mitigate depolarization and reduce proton leak through the mitochondrial ADP/ATP translocase ANT1.^91^ The tricationic peptide’s accumulation at the IMM’s surface could increase the local pH of the IMM interfacial region, thus preventing lipids from being protonated and inhibiting proton leak caused by lipid cycling via ANT1.^92–94^

On the other hand, our results on SS-31 inhibiting electroporation in CL-containing bilayers, reducing ion leak, and spontaneously transporting across the bilayer point toward an additional, functionally consistent mechanism for reducing proton leak. When Δ𝜇#_𝐻+_ is abnormally large, thermal fluctuations in ion distributions in the solvent could create a strong enough electric field and thus the right energetic conditions for spontaneous electroporation of the IMM. In that event, SS-31 could curb proton leak directly, by sterically occluding the pore’s opening and repelling cation flux electrostatically, and/or indirectly, by peptide transport through the pore across the bilayer. A representative example of this happening in our simulations is presented in **Fig. 6B-E**. SS-31 carries at +3 charge and its transmembrane transport would theoretically reduce the electrical drive for cations, like protons, to flow down their concentration gradient into the matrix.

Taken together, these results build on our previous work studying the potential membrane-mediated mechanism of MTPs in which we showed that MTPs increase *C_m_*.^14^ It also confirms that an increase in *C_m_* leads to a decrease in *ΔΨ_m_*, and offers mechanistic insight into experimental studies showing that SS-31 can prevent hyperpolarization of the IMM’s *ΔΨ_m_* in isolated mitochondria^12^ and inhibit overproduction of ROS in dysfunctional mitochondria.^16^ Under normal conditions, transient spikes in the of 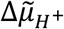 can occur (e.g., during state 4 respiration), which leads to ROS overproduction. Our results suggest that SS-31 increasing the *C_m_* could dilute the effect of spikes in the *ΔpH* by decreasing the electrical component of 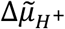 (*ΔΨ_m_*) such that even if the chemical component (*ΔpH*) goes up, there would be little to no net change in the 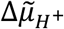 which in turn could curb ROS over-production. It also means that the IMM could support a larger 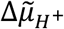 which may explain how MTPs accelerate H^+^-driven ATP production.

### Methodological limitations and correspondence to physiological conditions

The *ΔΨ_m_* values that triggered electroporation in our unbiased simulations are considerably higher than the ∼0.15 V reported across healthy IMM.^1^ In fact, the observed threshold for transient electroporation in CL-containing membranes (∼1.9 V) exceeds even those under pathological conditions.^10^ However, the values are consistent with previous MD studies measuring transmembrane potentials at similar charge imbalances.^29,31,44,95^ Importantly, experimental measurements of *ΔΨ_m_* represent spatial and temporal averages; they likely obscure short-lived or spatially localized peaks in membrane potential that could drive poration on sub-microscopic scales.

Recent studies using super resolution microscopy have revealed subtle heterogeneity in *ΔΨ_m_* both between and within cristae, and future methodological advances may uncover larger variations at finer spatial and temporal scales. Further, *ΔΨ_m_* -induced pore formation occurs is expected to be a self-limiting process, as ion flux will rapidly dissipate the potential. Thus, large-amplitude fluctuations in *ΔΨ_m_* could transiently exceed classical thresholds for electroporation but remain undetectable due to their short lifetime and subdiffraction spatial scale.

Another factor could be the presence of electrostatically localized proton domains in the IMM, which could locally raise the local electrochemical proton potential gradient near the IMM surface.^96^ Passive (a.k.a. unassisted) divalent cation transport (i.e., electroporation) has been shown to happen in giant unilamellar vesicles (GUVs) at much lower *ΔΨ_m_* values, with two studies using second harmonic microscopy reporting unassisted ion transport happening at 200-500 mV.^97,98^ The latter studies report observing local transient *ΔΨ_m_* fluctuations of several hundred millivolts randomly scattered across the surface of GUVs. The authors attribute the trigger for passive cation transport to these transient fluctuations and the downstream local effects they have on the bilayer and electric field. Further, they report timescales of passive divalent cation flux rates ranging from 20 to 200 μs per ion in GUVs depending on the lipid composition.

To evaluate whether our model captures these physiological behaviors, we extrapolated pore formation kinetics to lower *ΔΨ_m_* values using a voltage-dependent free energy model. Although classical electroporation theory predicts a quadratic dependence of the energy barrier on voltage,^99^ our data were better described by a model in which the energy barrier decreases linearly with *ΔΨ_m_*:

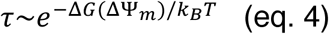

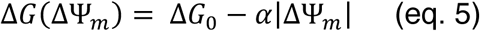

This model yielded a superior fit to the simulation data (r² = 0.9996) compared to a simple linear model (r² = 0.8441) (**Fig. S18**). The deviation from classical behavior may reflect system-specific factors — such as peptide–membrane interactions, bilayer deformation, or lipid heterogeneity — that alter the voltage-dependence of pore formation. Extrapolation of the model to low *ΔΨ_m_* values predicted pore formation timescales on the order of 10 µs, consistent with the experimentally measured passive ion transport rates in GUVs.^98^ This agreement serves as an important validation of the model, supporting its applicability not only under the high-voltage conditions of our simulations but also across more physiological voltage regimes.

These insights suggest that electroporation may occur even at lower voltages in large or long-lived systems. The larger the bilayer and the longer the observation window, the greater the probability that spontaneous, ion-driven poration events occur. Thus, while our simulations captured poration only at high voltages, similar processes may arise at physiological *ΔΨ_m_* in larger, more complex mitochondrial membranes on biological timescales.. Moreover, the difference in ion transport timescales between different lipid compositions is supportive of the difference we saw in electroporation propensity between our systems, with the 20% CL bilayer requiring a larger *δ_TM_* than the all-PC bilayer.

In summary, our results demonstrate that mitochondrial membrane composition and MTPs such as SS-31 can substantially influence the electrochemical conditions under which electroporation occurs. By increasing membrane capacitance and modulating structural integrity, CL and SS-31 act as key regulators of bilayer stability under electrochemical stress. These findings not only expand our mechanistic understanding of mitochondrial membrane biophysics but also offer new perspectives on how MTPs may protect against oxidative damage and support efficient energy production.

## Acknowledgements

This work was supported by the National Institutes of Health (Grant R35-GM119762 to E.R.M. and Grant R01-AG065879 to N.N.A.). The computational work performed on this project was done with help from the Storrs High-Performance Computing cluster. We would like to thank the UConn Storrs HPC and HPC team for providing the resources and support that contributed to these results.

