## Supporting Information for "Mitochondria-Targeted Peptides and Bilayer Composition Modulate Membrane Electroporation Under Elevated Electrochemical Stress"

Supporting information contains 18 supplementary figures, S1-S18.

**Double Bilayer 20:80 CL:PC System WITH peptides and NO ions swapped**

| | | | |
|---|---|---|---|
| | *Bilayer 1* | | |
| **IMS** | | Peptides | Cardiolipin |
| | | 10 | 30 |
| Compartmental Charge | | Sodium | Chloride |
| 0 | 100 mM NaCl | 43 | 43 |
| | Neutralizing | 30 | 0 |
| | | **73** | **43** |
| | *Bilayer 2* | | |
| **Matrix** | | Peptides | Cardiolipin |
| | | 0 | 30 |
| Compartmental Charge | | Sodium | Chloride |
| 0 | 100 mM NaCl | 43 | 43 |
| | Neutralizing | 60 | 0 |
| | | **103** | **43** |
| **System Charge** | **CIPL (IMS - MTX) / (75 lip/leaflet)** | | |
| 0 | 0 | | |

**Double Bilayer 20:80 CL:PC System with NO peptides and NO ions swapped**

| | | | |
|---|---|---|---|
| | *Bilayer 1* | | |
| **IMS** | | Peptides | Cardiolipin |
| | | 0 | 30 |
| Compartmental Charge | | Sodium | Chloride |
| 0 | 100 mM NaCl | 43 | 43 |
| | Neutralizing | 60 | 0 |
| | | **103** | **43** |
| | *Bilayer 2* | | |
| **Matrix** | | Peptides | Cardiolipin |
| | | 0 | 30 |
| Compartmental Charge | | Sodium | Chloride |
| 0 | 100 mM NaCl | 43 | 43 |
| | Neutralizing | 60 | 0 |
| | | **103** | **43** |
| **System Charge** | **CIPL (IMS - MTX) / (75 lip/leaflet)** | | |
| 0 | 0 | | |

**Double Bilayer 20:80 CL:PC System WITH peptides and ONE ion swapped**

| | | | |
|---|---|---|---|
| | *Bilayer 1* | | |
| **IMS** | | Peptides | Cardiolipin |
| | | 10 | 30 |
| Compartmental Charge | | Sodium | Chloride |
| 2 | 100 mM NaCl | 43 | 43 |
| | Neutralizing | 30 | 0 |
| | Ion Imbalance | **1** | **-1** |
| | | **74** | **42** |
| | *Bilayer 2* | | |
| **Matrix** | | Peptides | Cardiolipin |
| | | 0 | 30 |
| Compartmental Charge | | Sodium | Chloride |
| -2 | 100 mM NaCl | 43 | 43 |
| | Neutralizing | 60 | 0 |
| | Ion Imbalance | **-1** | **1** |
| | | **102** | **44** |
| **System Charge** | **CIPL (IMS - MTX) / (75 lip/leaflet)** | | |
| 0 | 0.05333333333 | | |

**Double Bilayer 20:80 CL:PC System with NO peptides and ONE ion swapped**

| | | | |
|---|---|---|---|
| | *Bilayer 1* | | |
| **IMS** | | Peptides | Cardiolipin |
| | | 0 | 30 |
| Compartmental Charge | | Sodium | Chloride |
| 2 | 100 mM NaCl | 43 | 43 |
| | Neutralizing | 60 | 0 |
| | Ion Imbalance | **1** | **-1** |
| | | **104** | **42** |
| | *Bilayer 2* | | |
| **Matrix** | | Peptides | Cardiolipin |
| | | 0 | 30 |
| Compartmental Charge | | Sodium | Chloride |
| -2 | 100 mM NaCl | 43 | 43 |
| | Neutralizing | 60 | 0 |
| | Ion Imbalance | **-1** | **1** |
| | | **102** | **44** |
| **System Charge** | **CIPL (IMS - MTX) / (75 lip/leaflet)** | | |
| 0 | 0.05333333333 | | |

**Double Bilayer 20:80 CL:PC System WITH peptides and TWO ions swapped**

| | | | |
|---|---|---|---|
| | *Bilayer 1* | | |
| **IMS** | | Peptides | Cardiolipin |
| | | 10 | 30 |
| Compartmental Charge | | Sodium | Chloride |
| 4 | 100 mM NaCl | 43 | 43 |
| | Neutralizing | 30 | 0 |
| | Ion Imbalance | **2** | **-2** |
| | | **75** | **41** |
| | *Bilayer 2* | | |
| **Matrix** | | Peptides | Cardiolipin |
| | | 0 | 30 |
| Compartmental Charge | | Sodium | Chloride |
| -4 | 100 mM NaCl | 43 | 43 |
| | Neutralizing | 60 | 0 |
| | Ion Imbalance | **-2** | **2** |
| | | **101** | **45** |
| **System Charge** | **CIPL (IMS - MTX) / (75 lip/leaflet)** | | |
| 0 | 0.1066666667 | | |

**Double Bilayer 20:80 CL:PC System with NO peptides and TWO ions swapped**

| | | | |
|---|---|---|---|
| | *Bilayer 1* | | |
| **IMS** | | Peptides | Cardiolipin |
| | | 0 | 30 |
| Compartmental Charge | | Sodium | Chloride |
| 4 | 100 mM NaCl | 43 | 43 |
| | Neutralizing | 60 | 0 |
| | Ion Imbalance | **2** | **-2** |
| | | **105** | **41** |
| | *Bilayer 2* | | |
| **Matrix** | | Peptides | Cardiolipin |
| | | 0 | 30 |
| Compartmental Charge | | Sodium | Chloride |
| -4 | 100 mM NaCl | 43 | 43 |
| | Neutralizing | 60 | 0 |
| | Ion Imbalance | **-2** | **2** |
| | | **101** | **45** |
| **System Charge** | **CIPL (IMS - MTX) / (75 lip/leaflet)** | | |
| 0 | 0.1066666667 | | |

### Supplementary figure 1 - Schematic representation of 20:80 TOCL: POPC bilayer systems with and without SS-31 (part 1)

Note that CIPL stands for charge imbalance per lipid, IMS stands for intermembrane space, and MTX stands for matrix.

**Double Bilayer 20:80 CL:PC System WITH peptides and THREE ions swapped**

| | | | |
|---|---|---|---|
| | | *Bilayer 1* | |
| **IMS** | | Peptides | Cardiolipin |
| | | 10 | 30 |
| Compartmental Charge | | Sodium | Chloride |
| 6 | 100 mM NaCl | 43 | 43 |
| | Neutralizing | 30 | 0 |
| | Ion Imbalance | **3** | **-3** |
| | | **76** | **40** |
| | | *Bilayer 2* | |
| **Matrix** | | Peptides | Cardiolipin |
| | | 0 | 30 |
| Compartmental Charge | | Sodium | Chloride |
| -6 | 100 mM NaCl | 43 | 43 |
| | Neutralizing | 60 | 0 |
| | Ion Imbalance | **-3** | **3** |
| | | **100** | **46** |
| **System Charge** | | **CIPL (IMS - MTX) / (75 lip/leaflet)** | |
| 0 | | 0.16 | |

**Double Bilayer 20:80 CL:PC System with NO peptides and THREE ions swapped**

| | | | |
|---|---|---|---|
| | | *Bilayer 1* | |
| **IMS** | | Peptides | Cardiolipin |
| | | 0 | 30 |
| Compartmental Charge | | Sodium | Chloride |
| 6 | 100 mM NaCl | 43 | 43 |
| | Neutralizing | 60 | 0 |
| | Ion Imbalance | **3** | **-3** |
| | | **106** | **40** |
| | | *Bilayer 2* | |
| **Matrix** | | Peptides | Cardiolipin |
| | | 0 | 30 |
| Compartmental Charge | | Sodium | Chloride |
| -6 | 100 mM NaCl | 43 | 43 |
| | Neutralizing | 60 | 0 |
| | Ion Imbalance | **-3** | **3** |
| | | **100** | **46** |
| **System Charge** | | **CIPL (IMS - MTX) / (75 lip/leaflet)** | |
| 0 | | 0.16 | |

**Double Bilayer 20:80 CL:PC System WITH peptides and FOUR ions swapped**

| | | | |
|---|---|---|---|
| | | *Bilayer 1* | |
| **IMS** | | Peptides | Cardiolipin |
| | | 10 | 30 |
| Compartmental Charge | | Sodium | Chloride |
| 8 | 100 mM NaCl | 43 | 43 |
| | Neutralizing | 30 | 0 |
| | Ion Imbalance | **4** | **-4** |
| | | **77** | **39** |
| | | *Bilayer 2* | |
| **Matrix** | | Peptides | Cardiolipin |
| | | 0 | 30 |
| Compartmental Charge | | Sodium | Chloride |
| -8 | 100 mM NaCl | 43 | 43 |
| | Neutralizing | 60 | 0 |
| | Ion Imbalance | **-4** | **4** |
| | | **99** | **47** |
| **System Charge** | | **CIPL (IMS - MTX) / (75 lip/leaflet)** | |
| 0 | | 0.2133333333 | |

**Double Bilayer 20:80 CL:PC System with NO peptides and FOUR ions swapped**

| | | | |
|---|---|---|---|
| | | *Bilayer 1* | |
| **IMS** | | Peptides | Cardiolipin |
| | | 0 | 30 |
| Compartmental Charge | | Sodium | Chloride |
| 8 | 100 mM NaCl | 43 | 43 |
| | Neutralizing | 60 | 0 |
| | Ion Imbalance | **4** | **-4** |
| | | **107** | **39** |
| | | *Bilayer 2* | |
| **Matrix** | | Peptides | Cardiolipin |
| | | 0 | 30 |
| Compartmental Charge | | Sodium | Chloride |
| -8 | 100 mM NaCl | 43 | 43 |
| | Neutralizing | 60 | 0 |
| | Ion Imbalance | **-4** | **4** |
| | | **99** | **47** |
| **System Charge** | | **CIPL (IMS - MTX) / (75 lip/leaflet)** | |
| 0 | | 0.2133333333 | |

### Supplementary figure 2 - Schematic representation of 20:80 TOCL: POPC bilayer systems with and without SS-31 (part 2)

Note that CIPL stands for charge imbalance per lipid, IMS stands for intermembrane space, and MTX stands for matrix.

**Double Bilayer PC System WITH peptides and NO ions swapped**

| | | | |
|---|---|---|---|
| | *Bilayer 1* | | |
| IMS | | Peptides | Cardiolipin |
| | | 10 | 0 |
| Compartmental Charge | | Sodium | Chloride |
| 0 | 100 mM NaCl | 42 | 42 |
| | Neutralizing | 0 | 30 |
| | | **42** | **72** |
| | *Bilayer 2* | | |
| Matrix | | Peptides | Cardiolipin |
| | | 0 | 0 |
| Compartmental Charge | | Sodium | Chloride |
| | 100 mM NaCl | 42 | 42 |
| 0 | Neutralizing | 0 | 0 |
| | | **42** | **42** |
| **System Charge** | | **CIPL (IMS - MTX) / (75 lip/leaflet)** | |
| 0 | | 0 | |

**Double Bilayer PC System with NO peptides and NO ions swapped**

| | | | |
|---|---|---|---|
| | *Bilayer 1* | | |
| IMS | | Peptides | Cardiolipin |
| | | 0 | 0 |
| Compartmental Charge | | Sodium | Chloride |
| | 100 mM NaCl | 42 | 42 |
| 0 | Ionic Equivalent | 30 | 0 |
| | Neutralizing | 0 | 30 |
| | | **72** | **72** |
| | *Bilayer 2* | | |
| Matrix | | Peptides | Cardiolipin |
| | | 0 | 0 |
| Compartmental Charge | | Sodium | Chloride |
| | 100 mM NaCl | 42 | 42 |
| 0 | Neutralizing | 0 | 0 |
| | | **42** | **42** |
| **System Charge** | | **CIPL (IMS - MTX) / (75 lip/leaflet)** | |
| 0 | | 0 | |

**Double Bilayer PC System WITH peptides and ONE ion swapped**

| | | | |
|---|---|---|---|
| | *Bilayer 1* | | |
| IMS | | Peptides | Cardiolipin |
| | | 10 | 0 |
| Compartmental Charge | | Sodium | Chloride |
| 2 | 100 mM NaCl | 42 | 42 |
| | Neutralizing | 0 | 30 |
| | Ion Imbalance | 1 | -1 |
| | | **43** | **71** |
| | *Bilayer 2* | | |
| Matrix | | Peptides | Cardiolipin |
| | | 0 | 0 |
| Compartmental Charge | | Sodium | Chloride |
| | 100 mM NaCl | 42 | 42 |
| -2 | Neutralizing | 0 | 0 |
| | Ion Imbalance | -1 | 1 |
| | | **41** | **43** |
| **System Charge** | | **CIPL (IMS - MTX) / (75 lip/leaflet)** | |
| 0 | | 0.05333333333 | |

**Double Bilayer PC System with NO peptides and ONE ion swapped**

| | | | |
|---|---|---|---|
| | *Bilayer 1* | | |
| IMS | | Peptides | Cardiolipin |
| | | 0 | 0 |
| Compartmental Charge | | Sodium | Chloride |
| | 100 mM NaCl | 42 | 42 |
| 2 | Ionic Equivalent | 30 | 0 |
| | Neutralizing | 0 | 30 |
| | Ion Imbalance | 1 | -1 |
| | | **73** | **71** |
| | *Bilayer 2* | | |
| Matrix | | Peptides | Cardiolipin |
| | | 0 | 0 |
| Compartmental Charge | | Sodium | Chloride |
| | 100 mM NaCl | 42 | 42 |
| -2 | Neutralizing | 0 | 0 |
| | Ion Imbalance | -1 | 1 |
| | | **41** | **43** |
| **System Charge** | | **CIPL (IMS - MTX) / (75 lip/leaflet)** | |
| 0 | | 0.05333333333 | |

**Double Bilayer PC System WITH peptides and TWO ions swapped**

| | | | |
|---|---|---|---|
| | *Bilayer 1* | | |
| IMS | | Peptides | Cardiolipin |
| | | 10 | 0 |
| Compartmental Charge | | Sodium | Chloride |
| 4 | 100 mM NaCl | 42 | 42 |
| | Neutralizing | 0 | 30 |
| | Ion Imbalance | 2 | -2 |
| | | **44** | **70** |
| | *Bilayer 2* | | |
| Matrix | | Peptides | Cardiolipin |
| | | 0 | 0 |
| Compartmental Charge | | Sodium | Chloride |
| | 100 mM NaCl | 42 | 42 |
| -4 | Neutralizing | 0 | 0 |
| | Ion Imbalance | -2 | 2 |
| | | **40** | **44** |
| **System Charge** | | **CIPL (IMS - MTX) / (75 lip/leaflet)** | |
| 0 | | 0.1066666667 | |

**Double Bilayer PC System with NO peptides and TWO ions swapped**

| | | | |
|---|---|---|---|
| | *Bilayer 1* | | |
| IMS | | Peptides | Cardiolipin |
| | | 0 | 0 |
| Compartmental Charge | | Sodium | Chloride |
| | 100 mM NaCl | 42 | 42 |
| 4 | Ionic Equivalent | 30 | 0 |
| | Neutralizing | 0 | 30 |
| | Ion Imbalance | 2 | -2 |
| | | **74** | **70** |
| | *Bilayer 2* | | |
| Matrix | | Peptides | Cardiolipin |
| | | 0 | 0 |
| Compartmental Charge | | Sodium | Chloride |
| | 100 mM NaCl | 42 | 42 |
| -4 | Neutralizing | 0 | 0 |
| | Ion Imbalance | -2 | 2 |
| | | **40** | **44** |
| **System Charge** | | **CIPL (IMS - MTX) / (75 lip/leaflet)** | |
| 0 | | 0.1066666667 | |

### Supplementary figure 3 - Schematic representation of 100% POPC bilayer systems with and without SS-31 (part 1)

Note that CIPL stands for charge imbalance per lipid, IMS stands for intermembrane space, and MTX stands for matrix.

**Double Bilayer PC System WITH peptides and THREE ions swapped**

| | | | |
|---|---|---|---|
| | *Bilayer 1* | | |
| **IMS** | | Peptides | Cardiolipin |
| | | 10 | 0 |
| Compartmental Charge | | Sodium | Chloride |
| 6 | 100 mM NaCl | 42 | 42 |
| | Neutralizing | 0 | 30 |
| | Ion Imbalance | 3 | -3 |
| | | **45** | **69** |
| | *Bilayer 2* | | |
| **Matrix** | | Peptides | Cardiolipin |
| | | 0 | 0 |
| Compartmental Charge | | Sodium | Chloride |
| | 100 mM NaCl | 42 | 42 |
| -6 | Neutralizing | 0 | 0 |
| | Ion Imbalance | -3 | 3 |
| | | **39** | **45** |
| **System Charge** | **CIPL (IMS - MTX) / (75 lip/leaflet)** | | |
| 0 | 0.16 | | |

**Double Bilayer PC System with NO peptides and THREE ions swapped**

| | | | |
|---|---|---|---|
| | *Bilayer 1* | | |
| **IMS** | | Peptides | Cardiolipin |
| | | 0 | 0 |
| Compartmental Charge | | Sodium | Chloride |
| | 100 mM NaCl | 42 | 42 |
| | Ionic Equivalent | 30 | 0 |
| 6 | Neutralizing | 0 | 30 |
| | Ion Imbalance | 3 | -3 |
| | | **75** | **69** |
| | *Bilayer 2* | | |
| **Matrix** | | Peptides | Cardiolipin |
| | | 0 | 0 |
| Compartmental Charge | | Sodium | Chloride |
| | 100 mM NaCl | 42 | 42 |
| -6 | Neutralizing | 0 | 0 |
| | Ion Imbalance | -3 | 3 |
| | | **39** | **45** |
| **System Charge** | **CIPL (IMS - MTX) / (75 lip/leaflet)** | | |
| 0 | 0.16 | | |

**Double Bilayer PC System WITH peptides and FOUR ions swapped**

| | | | |
|---|---|---|---|
| | *Bilayer 1* | | |
| **IMS** | | Peptides | Cardiolipin |
| | | 10 | 0 |
| Compartmental Charge | | Sodium | Chloride |
| 8 | 100 mM NaCl | 42 | 42 |
| | Neutralizing | 0 | 30 |
| | Ion Imbalance | 4 | -4 |
| | | **46** | **68** |
| | *Bilayer 2* | | |
| **Matrix** | | Peptides | Cardiolipin |
| | | 0 | 0 |
| Compartmental Charge | | Sodium | Chloride |
| | 100 mM NaCl | 42 | 42 |
| -8 | Neutralizing | 0 | 0 |
| | Ion Imbalance | -4 | 4 |
| | | **38** | **46** |
| **System Charge** | **CIPL (IMS - MTX) / (75 lip/leaflet)** | | |
| 0 | 0.2133333333 | | |

**Double Bilayer PC System with NO peptides and FOUR ions swapped**

| | | | |
|---|---|---|---|
| | *Bilayer 1* | | |
| **IMS** | | Peptides | Cardiolipin |
| | | 0 | 0 |
| Compartmental Charge | | Sodium | Chloride |
| | 100 mM NaCl | 42 | 42 |
| | Ionic Equivalent | 30 | 0 |
| 8 | Neutralizing | 0 | 30 |
| | Ion Imbalance | 4 | -4 |
| | | **76** | **68** |
| | *Bilayer 2* | | |
| **Matrix** | | Peptides | Cardiolipin |
| | | 0 | 0 |
| Compartmental Charge | | Sodium | Chloride |
| | 100 mM NaCl | 42 | 42 |
| -8 | Neutralizing | 0 | 0 |
| | Ion Imbalance | -4 | 4 |
| | | **38** | **46** |
| **System Charge** | **CIPL (IMS - MTX) / (75 lip/leaflet)** | | |
| 0 | 0.2133333333 | | |

### Supplementary figure 4 - Schematic representation of 100% POPC bilayer systems with and without SS-31 (part 2)

Note that CIPL stands for charge imbalance per lipid, IMS stands for intermembrane space, and MTX stands for matrix.

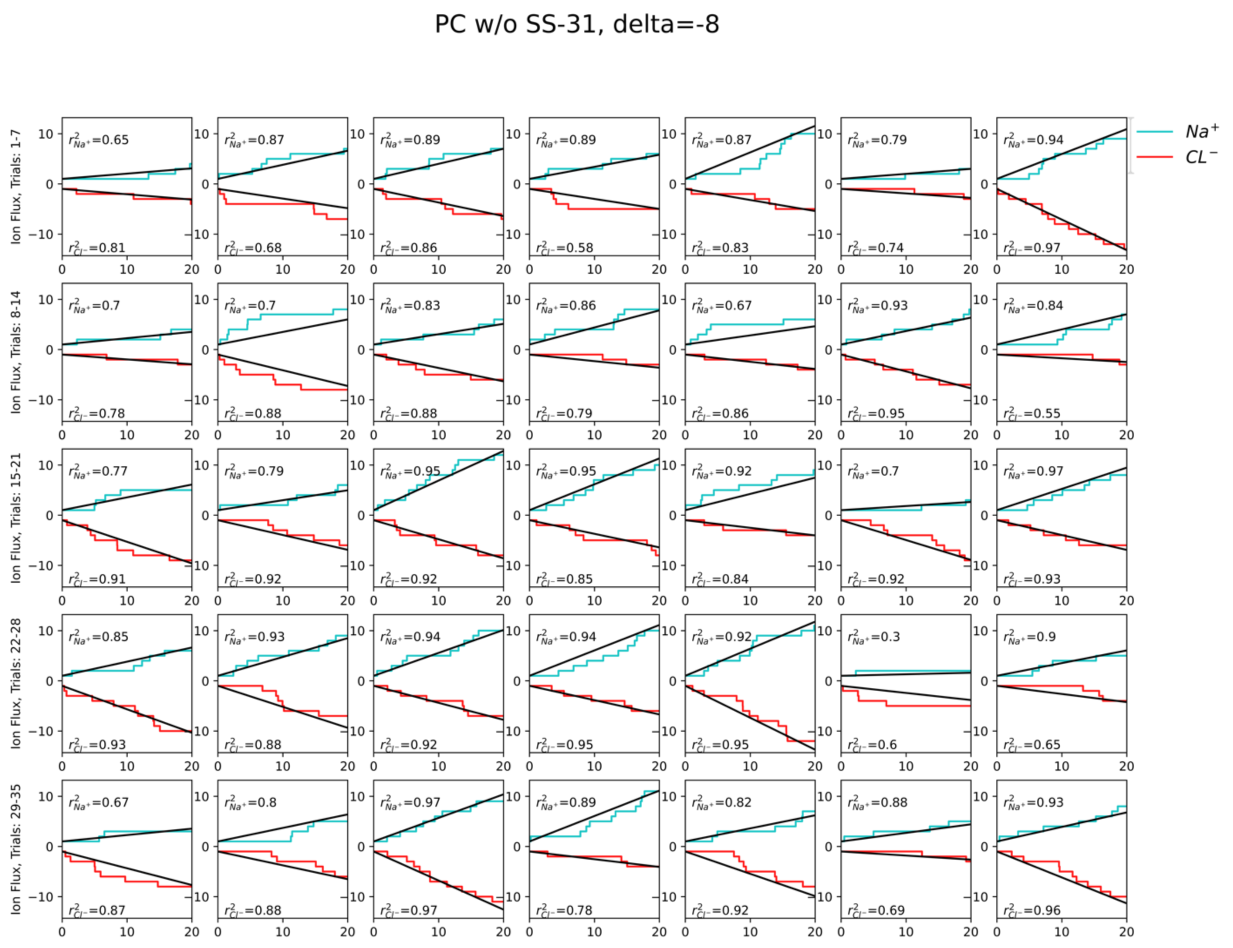


**Supplementary Figure 5 - Linear regression of raw ion flux from CompEL simulations of Apo all POPC bilayer at $\delta_{TM}$=-8**

Note that the y-axis is time in ns and that $r^2$ values are listed as N/A in a small number of trials whose slopes (ion flux rates) were not calculated by linear regression but were instead calculated manually due to very low ion flux rates. For more details, please see the methods.

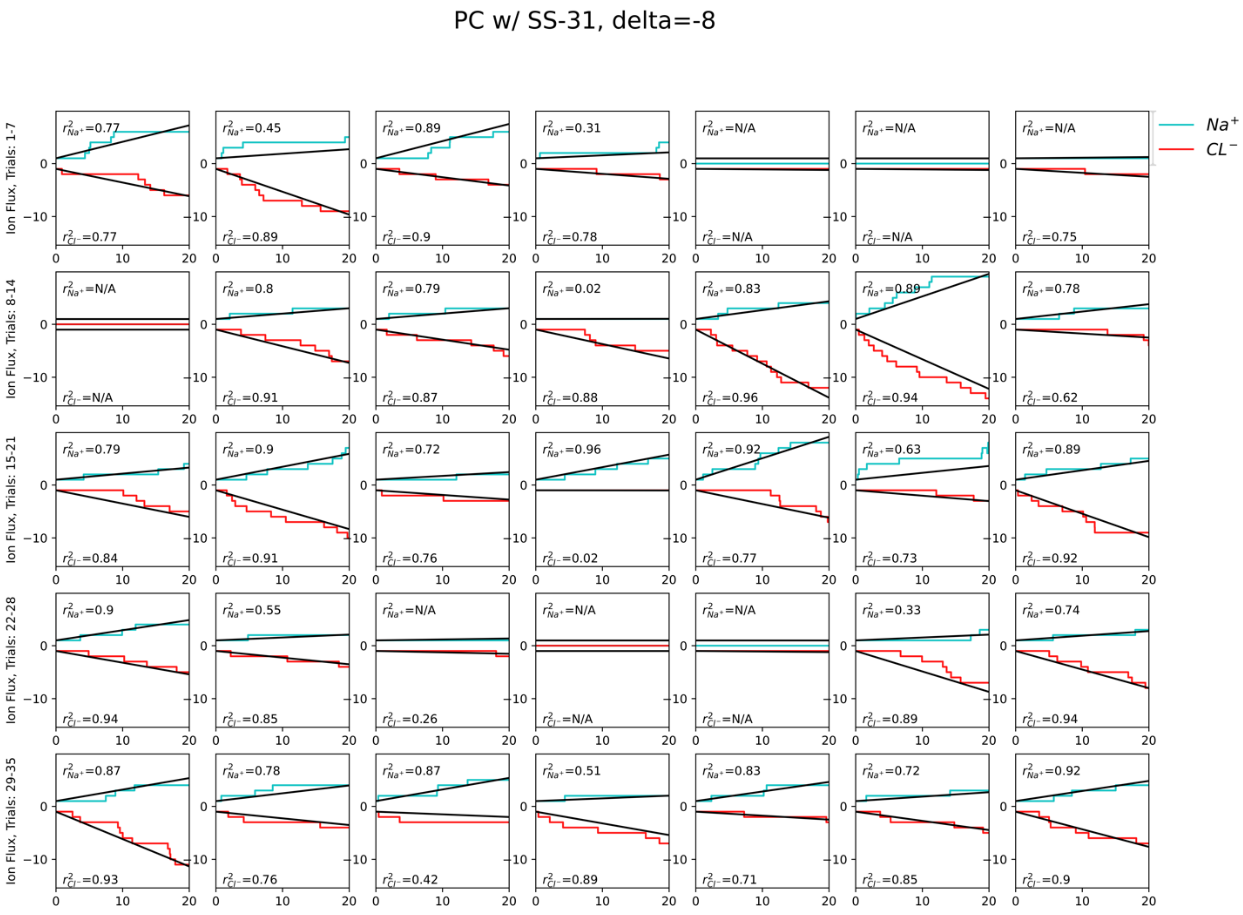


**Supplementary Figure 6 - Linear regression of raw ion flux from CompEL simulations of an SS-31-containing all POPC bilayer at $\delta_{TM}$=-8**

Note that the y-axis is time in ns and that $r^2$ values are listed as N/A in a small number of trials whose slopes (ion flux rates) were not calculated by linear regression but were instead calculated manually due to very low ion flux rates. For more details, please see the methods. In this system, there were two trials which exhibited no ion flux for either ion (trials 8 and 25). These trials were omitted from ion flux analyses.

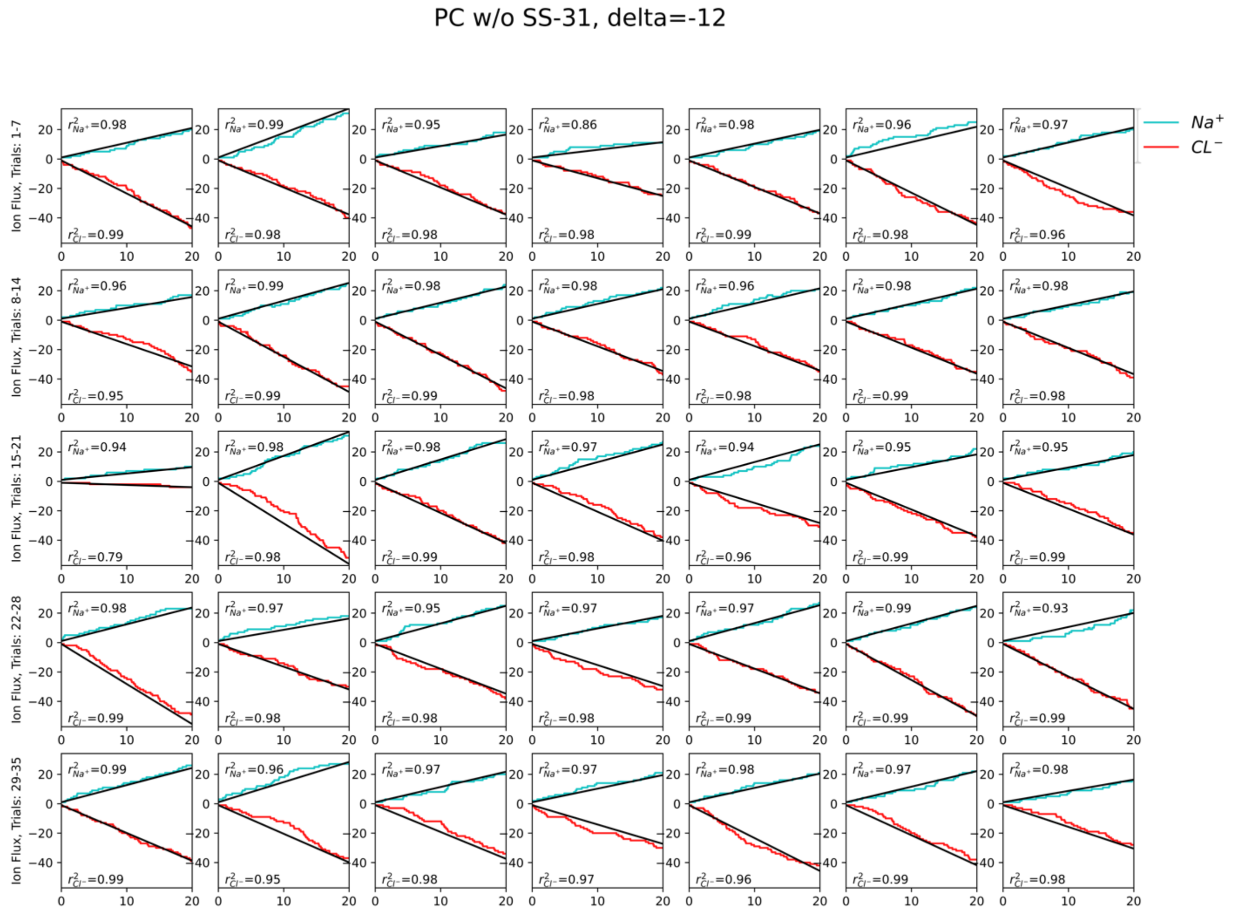


**Supplementary Figure 7 - Linear regression of raw ion flux from CompEL simulations of an Apo all POPC bilayer at $\delta_{TM}$=-12**

Note that the y-axis is time in ns and that $r^2$ values are listed as N/A in a small number of trials whose slopes (ion flux rates) were not calculated by linear regression but were instead calculated manually due to very low ion flux rates. For more details, please see the methods.

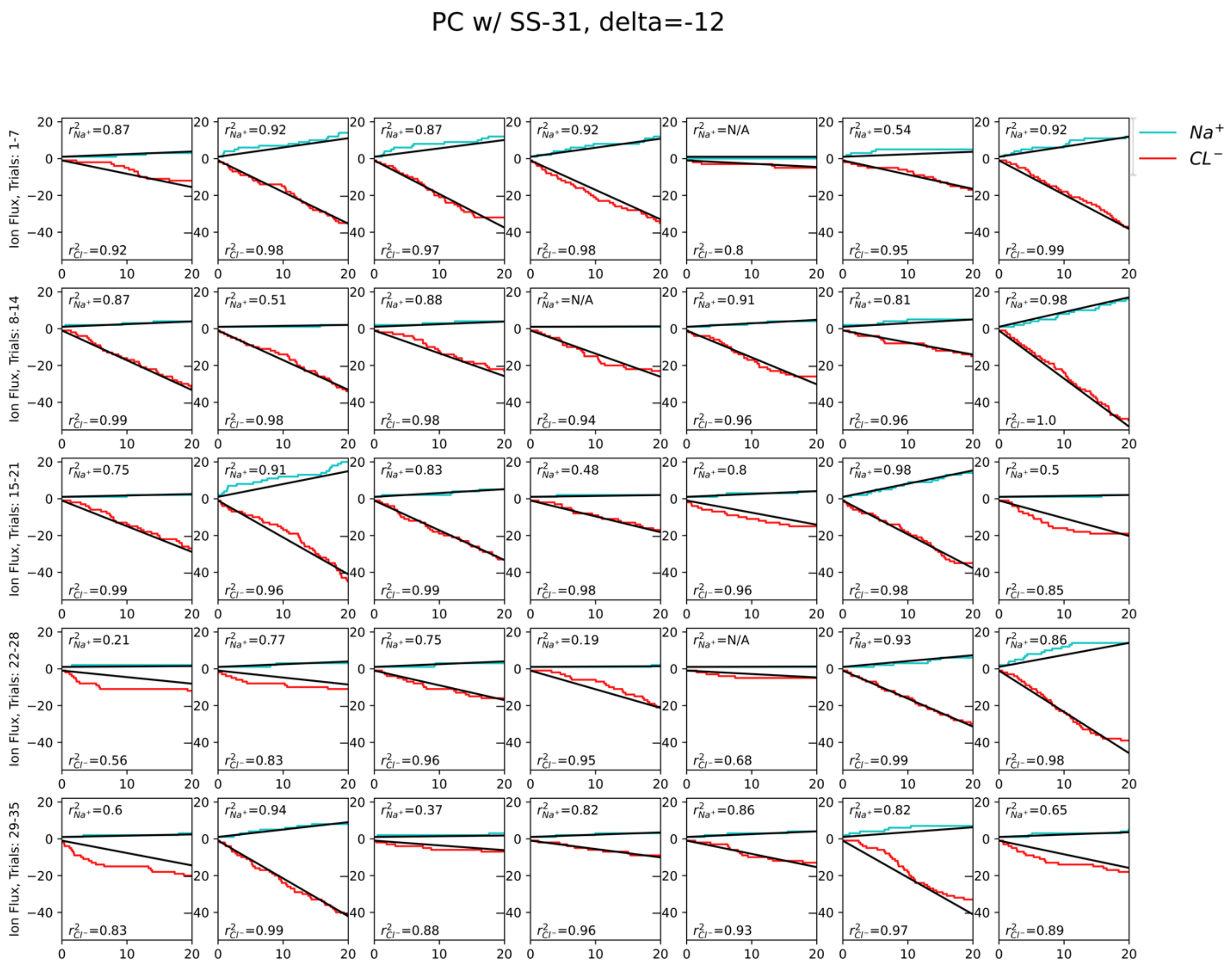


**Supplementary Figure 8 - Linear regression of raw ion flux from CompEL simulations of an SS-31-containing all POPC bilayer at $\delta_{TM}$=-12**

Note that the y-axis is time in ns and that $r^2$ values are listed as N/A in a small number of trials whose slopes (ion flux rates) were not calculated by linear regression but were instead calculated manually due to very low ion flux rates. For more details, please see the methods.

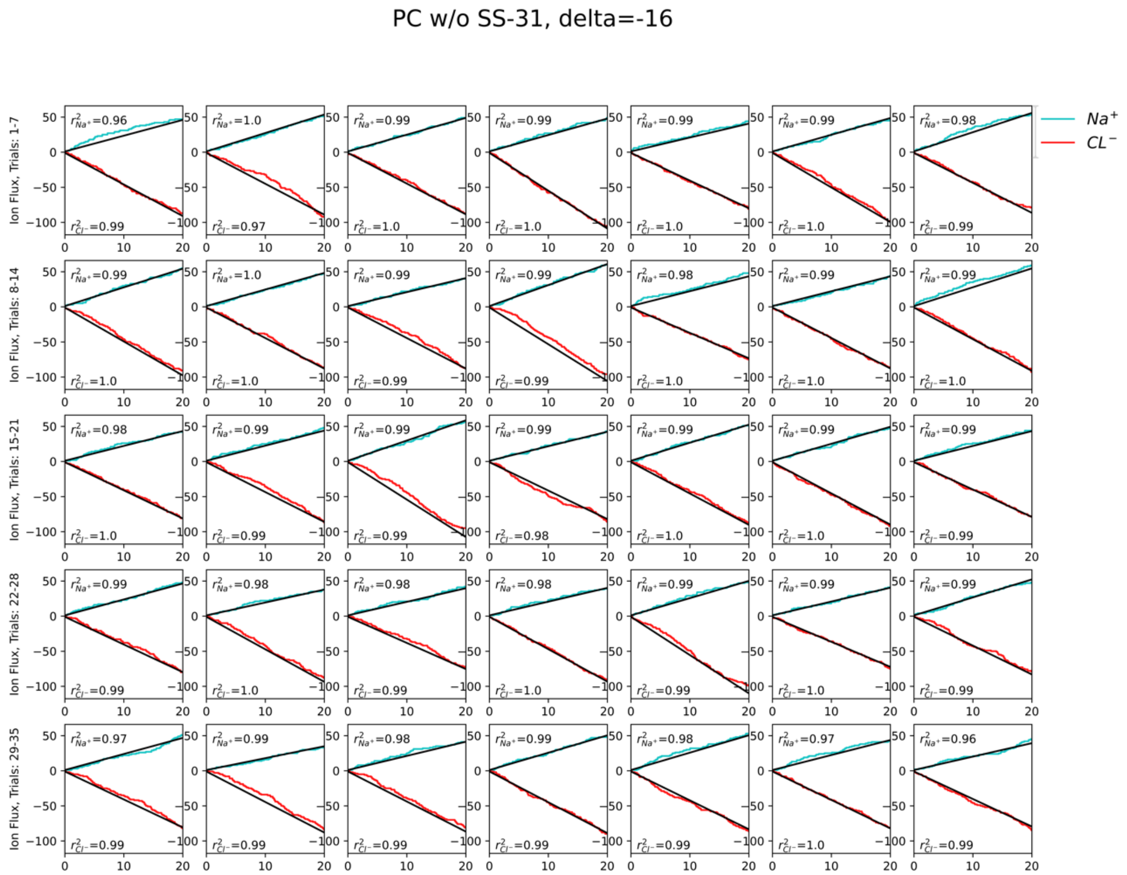


**Supplementary Figure 9 - Linear regression of raw ion flux from CompEL simulations of an Apo all POPC bilayer at $\delta_{TM}$=-16**

Note that the y-axis is time in ns and that $r^2$ values are listed as N/A in a small number of trials whose slopes (ion flux rates) were not calculated by linear regression but were instead calculated manually due to very low ion flux rates. For more details, please see the methods.

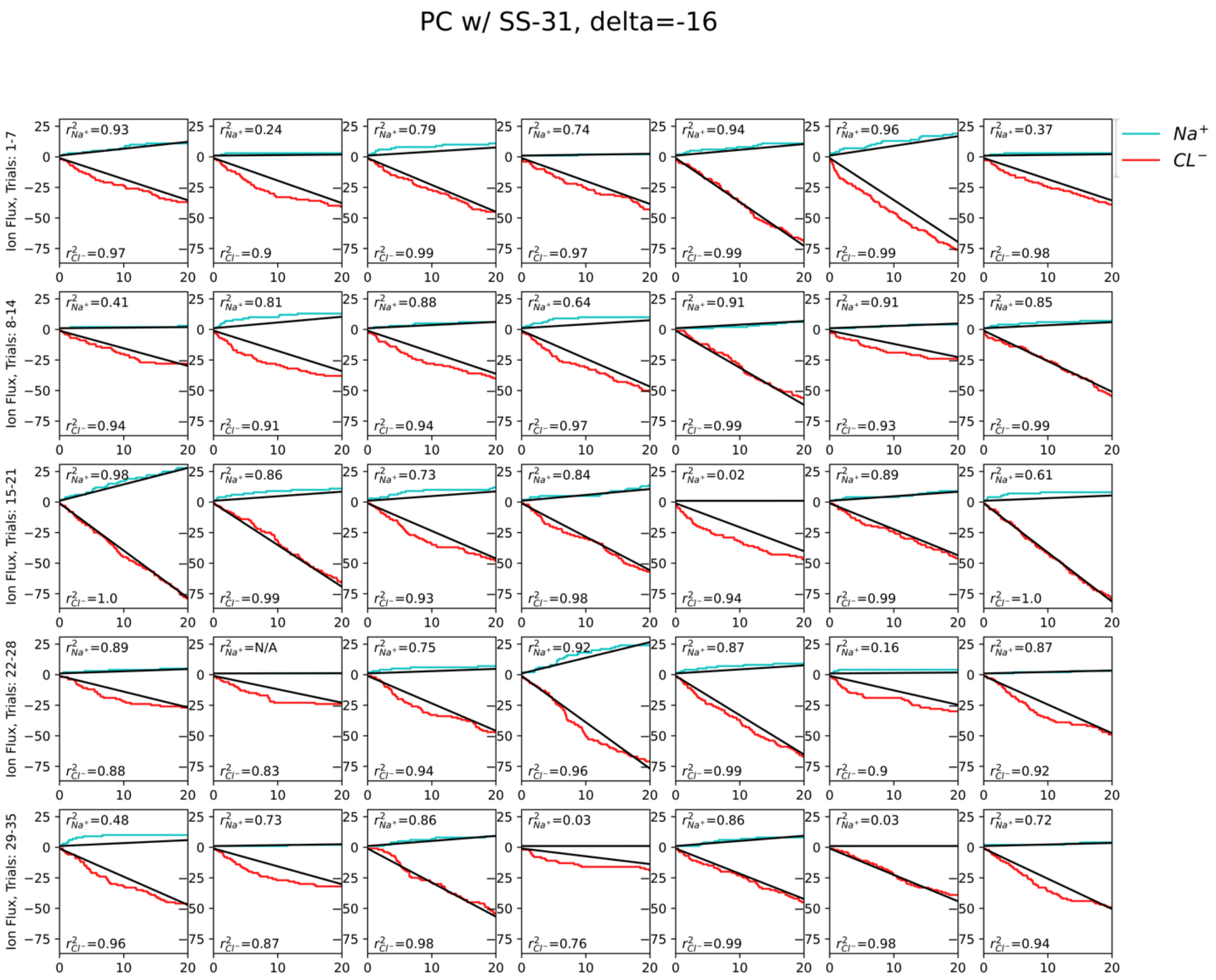


**Supplementary Figure 10 - Linear regression of raw ion flux from CompEL simulations of an SS-31-containing all POPC bilayer at $\delta_{TM}$=-16**

Note that the y-axis is time in ns and that $r^2$ values are listed as N/A in a small number of trials whose slopes (ion flux rates) were not calculated by linear regression but were instead calculated manually due to very low ion flux rates. For more details, please see the methods.

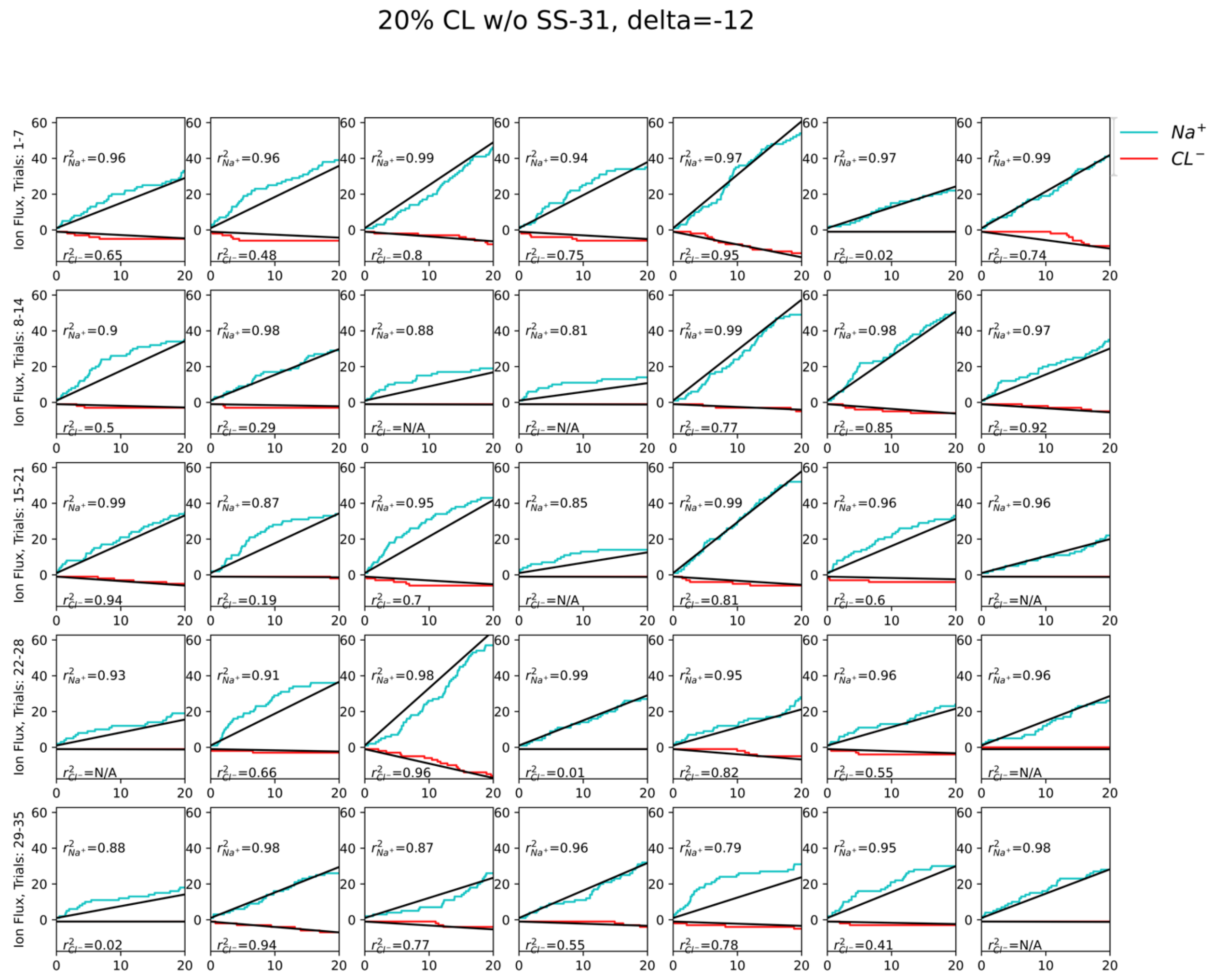


**Supplementary Figure 11 - Linear regression of raw ion flux from CompEL simulations of an Apo 20% CL bilayer at $\delta_{TM}$=-12**

Note that the y-axis is time in ns and that $r^2$ values are listed as N/A in a small number of trials whose slopes (ion flux rates) were not calculated by linear regression but were instead calculated manually due to very low ion flux rates. For more details, please see the methods.

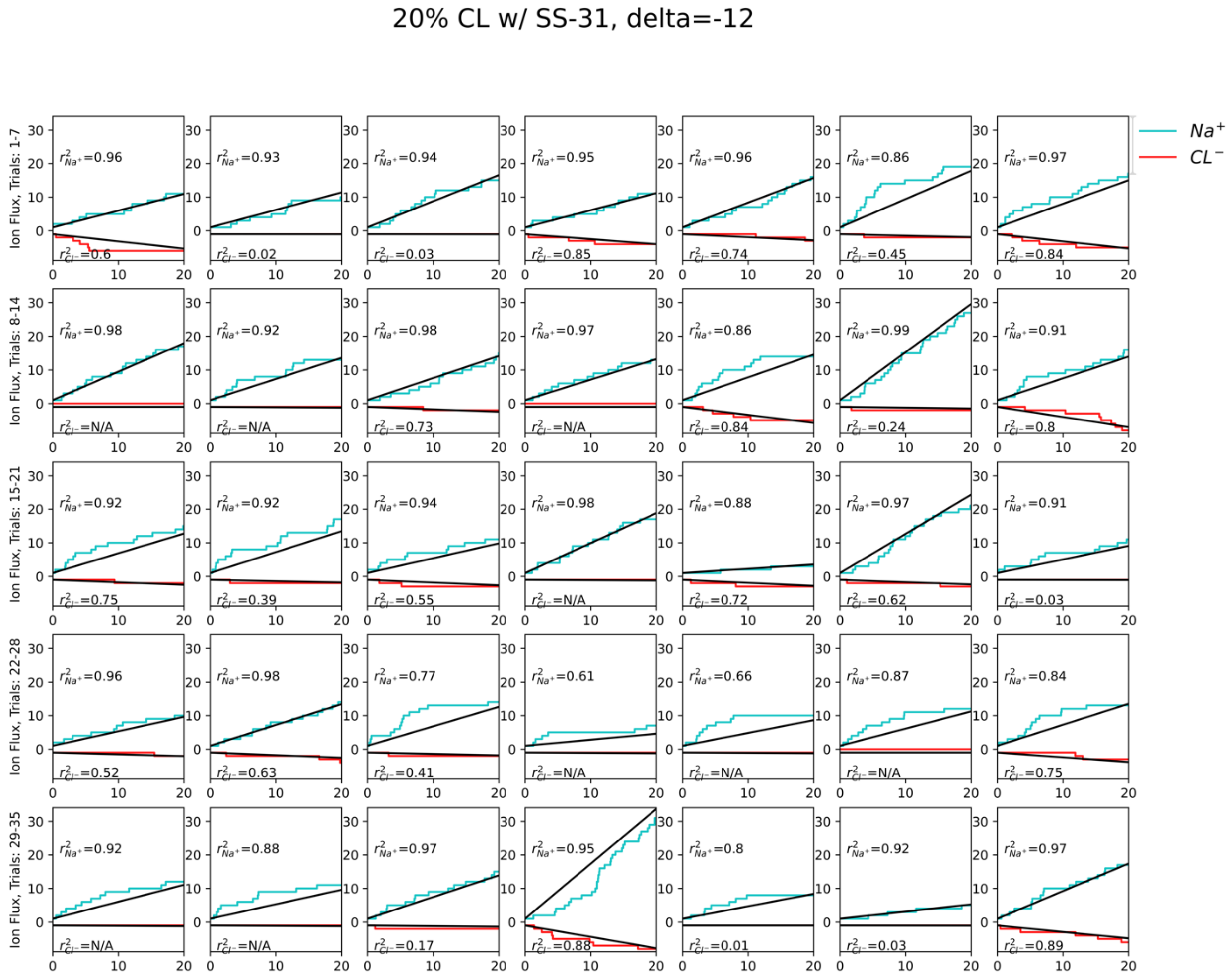


**Supplementary Figure 12 - Linear regression of raw ion flux from CompEL simulations of an SS-31-containing 20% CL bilayer at $\delta_{TM}$=-12**

Note that the y-axis is time in ns and that $r^2$ values are listed as N/A in a small number of trials whose slopes (ion flux rates) were not calculated by linear regression but were instead calculated manually due to very low ion flux rates. For more details, please see the methods.

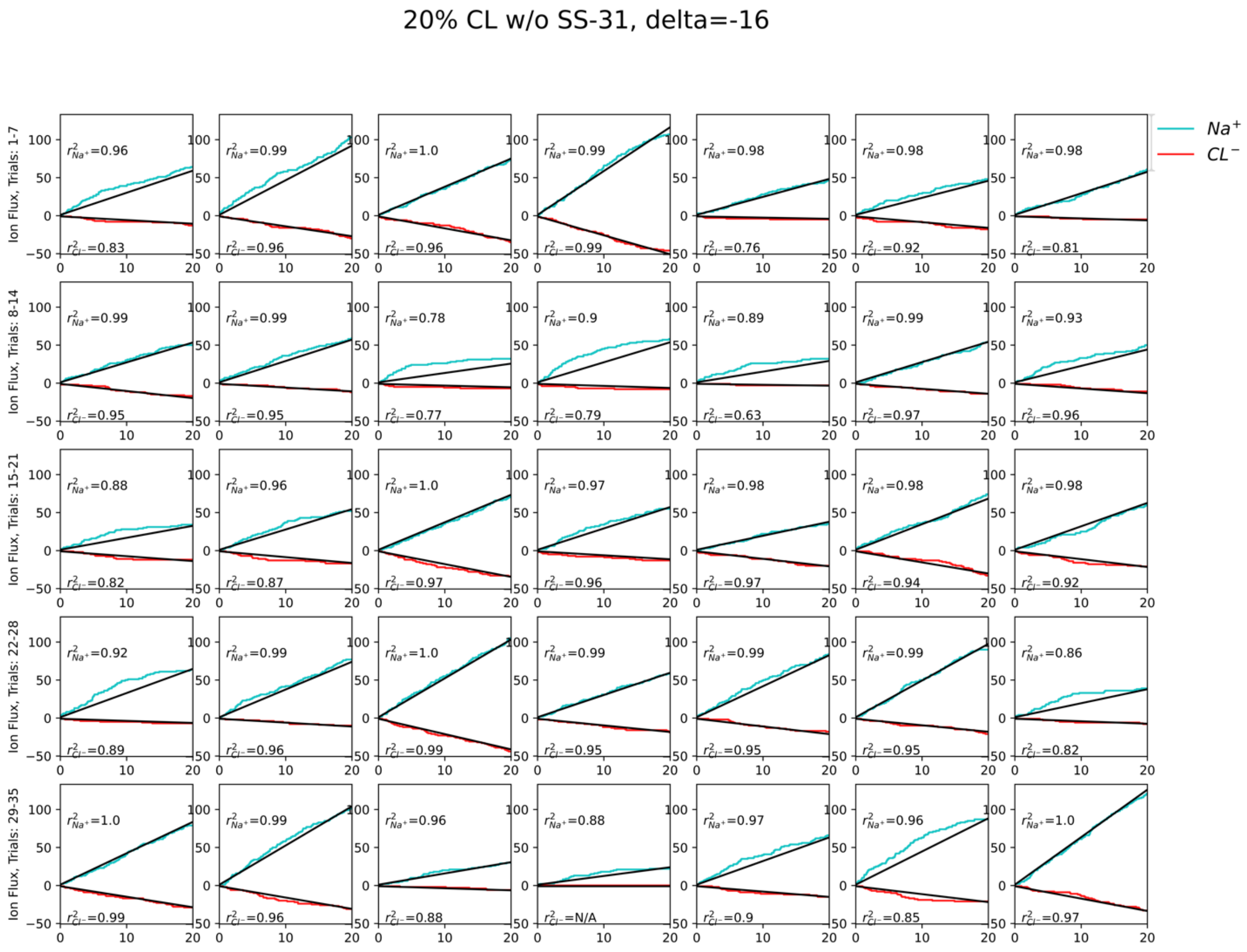


**Supplementary Figure 13 - Linear regression of raw ion flux from CompEL simulations of an Apo 20% CL bi-layer at $\delta_{TM}$=-16**

Note that the y-axis is time in ns and that $r^2$ values are listed as N/A in a small number of trials whose slopes (ion flux rates) were not calculated by linear regression but were instead calculated manually due to very low ion flux rates. For more details, please see the methods.

20% CL w/ SS-31, delta=-16

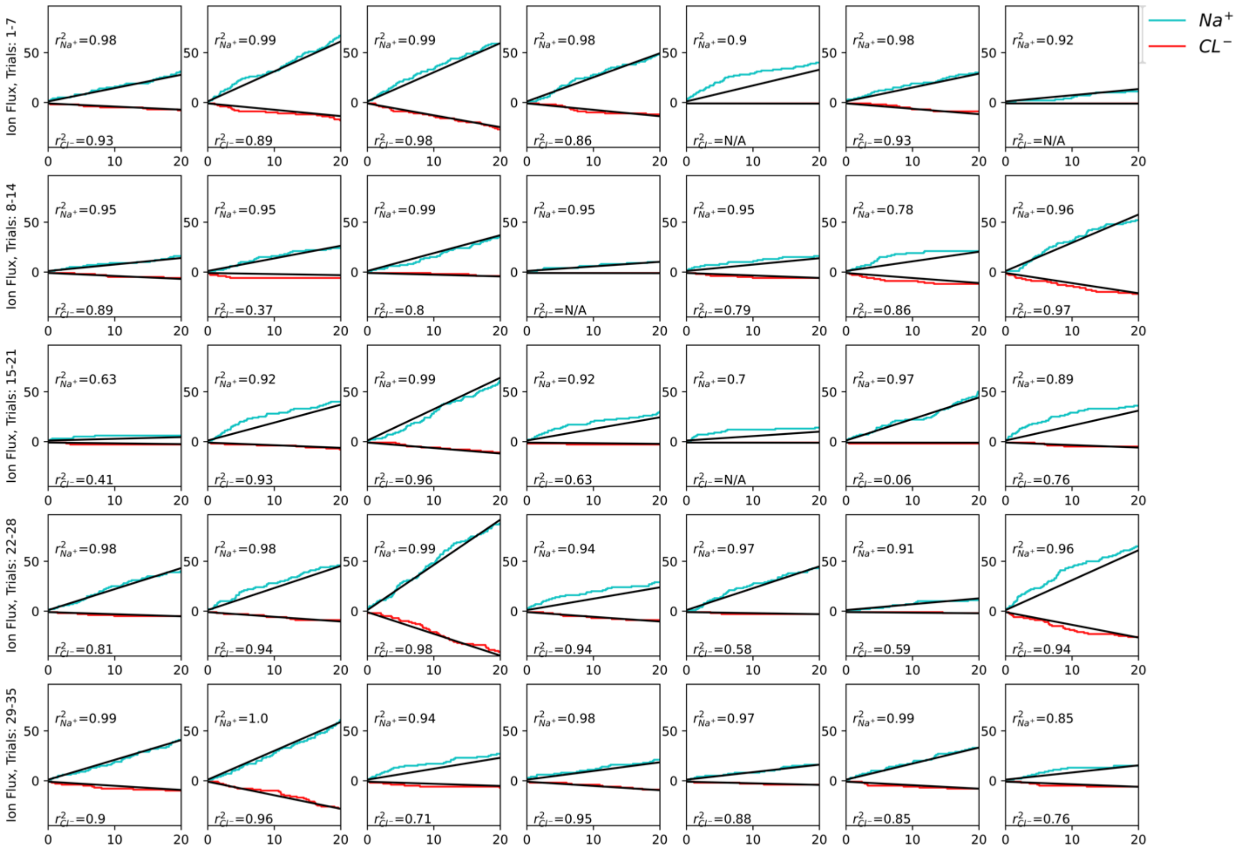

**Supplementary Figure 14 - Linear regression of raw ion flux from CompEL simulations of an SS-31-containing 20% CL bilayer at $\delta_{TM}$=-16**

Note that the y-axis is time in ns and that $r^2$ values are listed as N/A in a small number of trials whose slopes (ion flux rates) were not calculated by linear regression but were instead calculated manually due to very low ion flux rates. For more details, please see the methods.

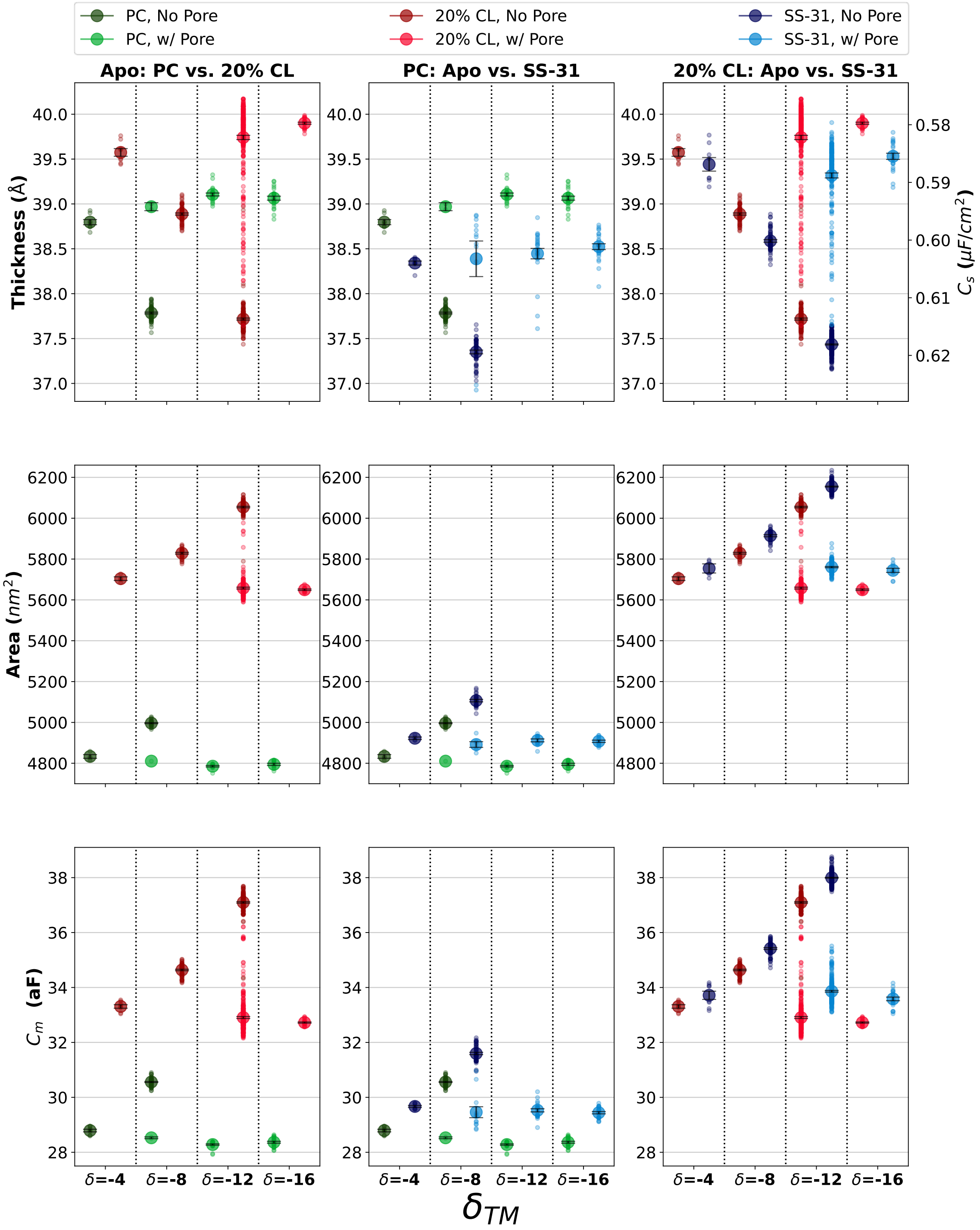


**Supplementary Figure 15. Comparison of Bilayer Properties and Total Capacitance Across Conditions**

Comparison of the average bilayer thickness (top row, left axis) and specific capacitance (top row, right axis), membrane surface area (middle row), and total capacitance (bottom row) among the conditions. The left column displays a comparison between the apo all POPC and apo 20% CL conditions; the middle column, apo vs. SS-31 in the all POPC bilayer; and the right column, apo vs. SS-31 in the 20% CL bilayer.

The small, color-coded circles represent the averages from each bilayer in individual simulations (see **Fig.** for n in each condition), and the larger circles with errors represents the means ± the standard errors. Note that the bilayer thickness is quantified as the distance between lipid phosphates in the upper and lower leaflets in each bilayer and that the y-axis for the capacitance is inverted. Also note that all results come from the end of simulations. The lighter colors represent systems which have electroporated, meaning the $\boldsymbol{\delta}_{TM}$ has changed from the initial simulation state shown in the X-axis label below each set of data.

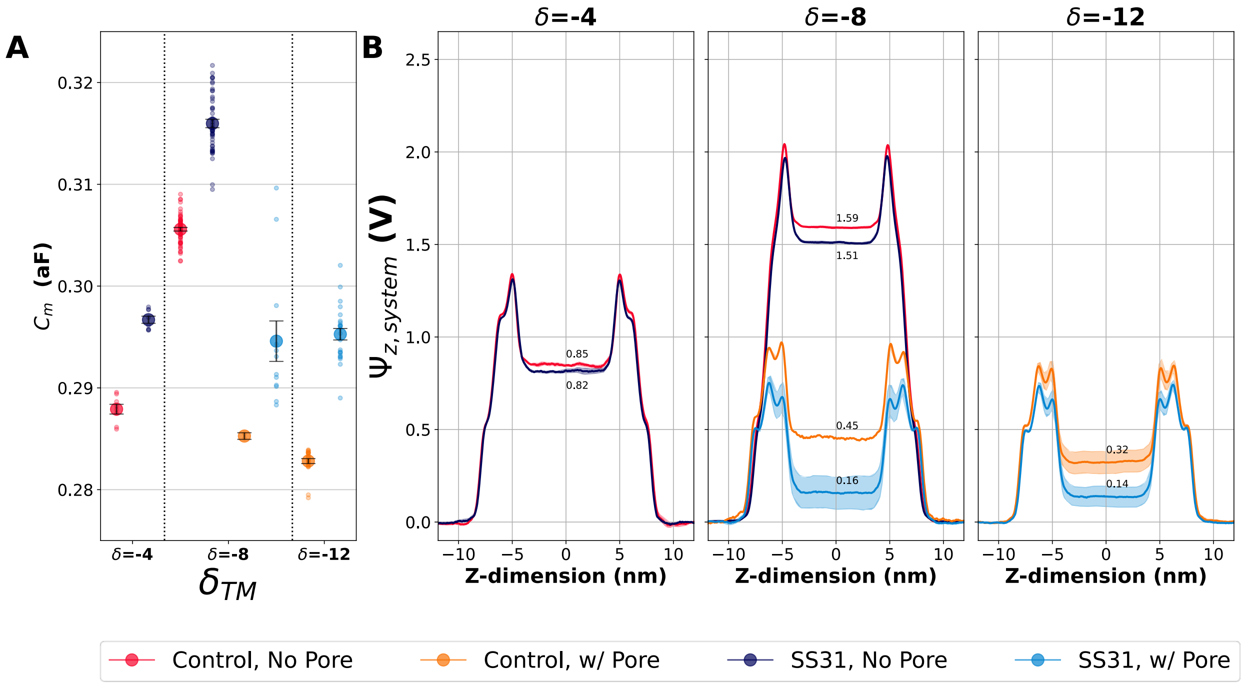


**Supplementary Figure 16. Electrostatic potential and membrane properties in all POPC bilayers with and without SS-31**

(A) Comparison of electrostatic potential between the apo all POPC bilayer and the SS-31-containing systems at different levels of exposure, as well as with and without unassisted ion leakage. (B) Comparison of the average bilayer thickness (left axis) and capacitance (right axis) among the conditions. The small, color-coded circles represent the averages from each bilayer in individual simulations (see **Fig. 1D**'s all POPC subsection for n in each condition), and the larger circles with errors represents the means ± the standard errors. Note that the bilayer thickness is quantified as the distance between lipid phosphates in the upper and lower leaflets in each bilayer and that the y-axis for the capacitance is inverted.

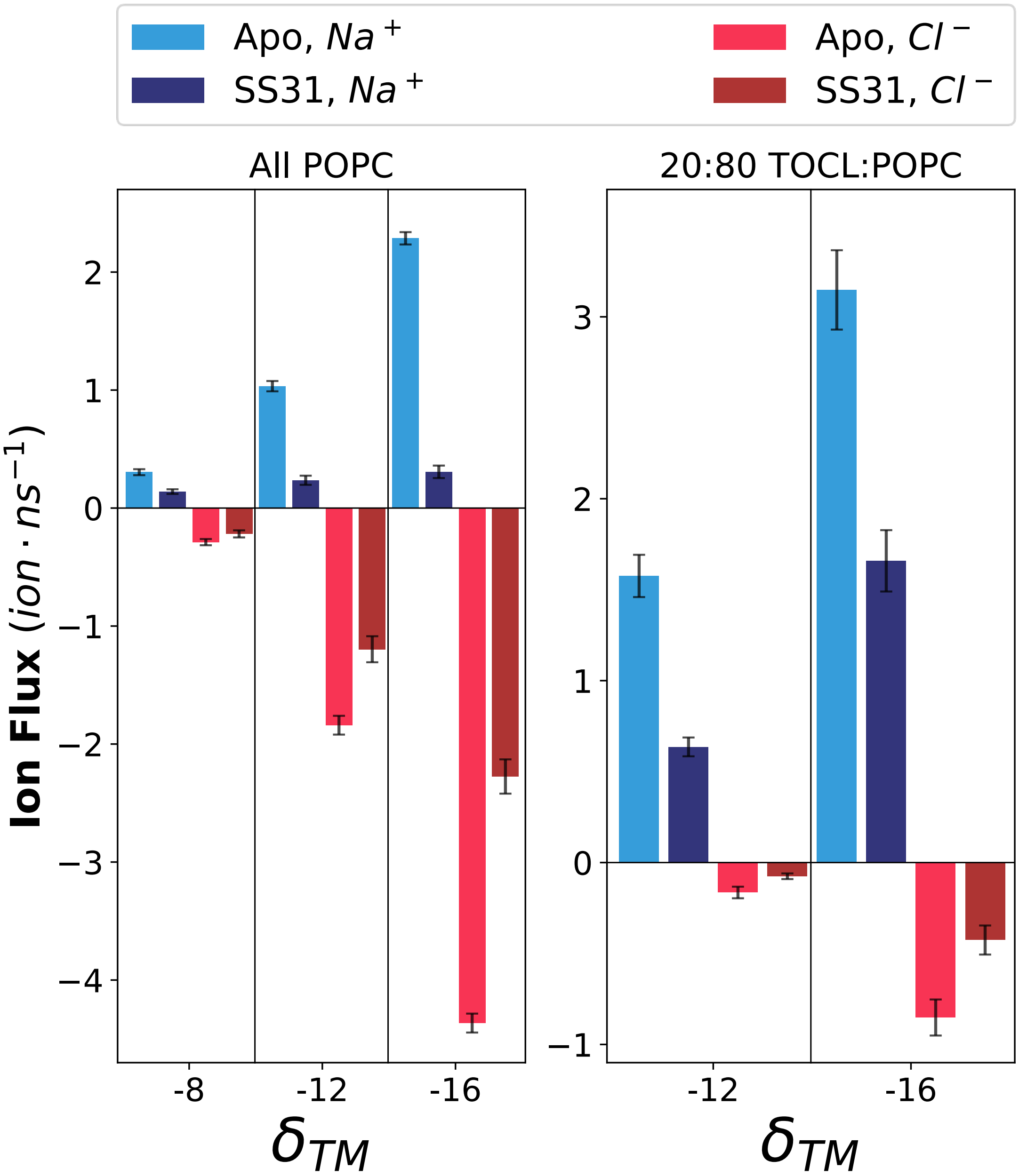


**Supplementary Figure 17. Comparison of raw ion flux rates from CompEL simulations**

Cation and anion flux rates were measured under different ion imbalances and for pure POPC (left) and 80:20 POPC:TOCL membranes (right).

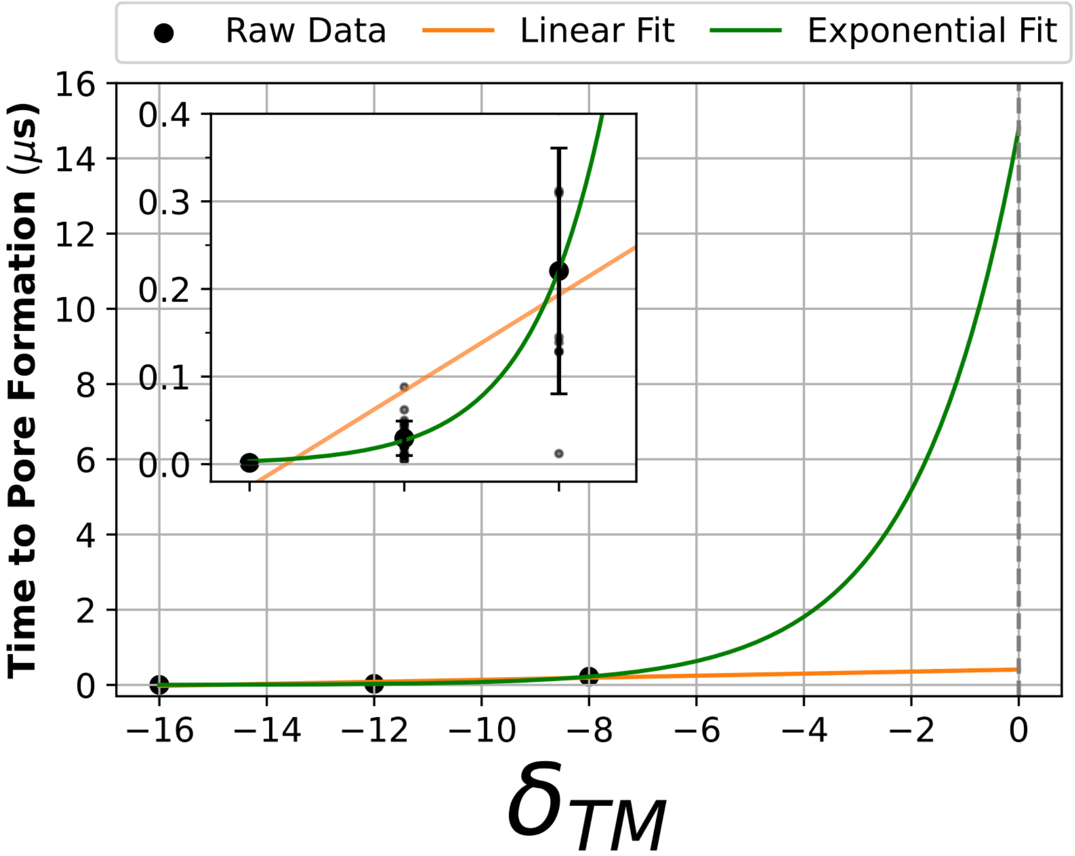


**Supplementary Figure 18. Fitting of pore formation kinetic data.**

Focused comparison of the amount of time before pore formation for the all PC bilayer systems. Data was pooled from the simulations with and without SS-31. Large black dots represent the average time to pore formation for a given $\delta_{TM}$ while the smaller black dots represent individual trials. Error bars are the standard deviation. The raw data were fit to both a linear and exponential curve to predict how the relationship between $\delta_{TM}$ and time to pore formation may behave at lower $\delta_{TM}$ values.
